# A Fluorescent Split Aptamer for Visualizing RNA-RNA Assembly *In Vivo*

**DOI:** 10.1101/109306

**Authors:** Khalid K. Alam, Kwaku D. Tawiah, Matthew F. Lichte, David Porciani, Donald H. Burke

## Abstract

RNA-RNA assembly governs key biological processes and is a powerful tool for engineering synthetic genetic circuits. Characterizing RNA assembly in living cells often involves monitoring fluorescent reporter proteins, which are at best indirect measures of underlying RNA-RNA hybridization events and are subject to additional temporal and load constraints associated with translation and activation of reporter proteins. In contrast, RNA aptamers that sequester small molecule dyes and activate their fluorescence are increasingly utilized in genetically-encoded strategies to report on RNA-level events. Split-aptamer systems have been rationally designed to generate signal upon hybridization of two or more discrete RNA transcripts, but none directly function when expressed *in vivo*. We reasoned that the improved physiological properties of the Broccoli aptamer enable construction of a split-aptamer system that could function in living cells. Here we present the Split-Broccoli system, in which self-assembly is nucleated by a thermostable, three-way junction RNA architecture and fluorescence activation requires both strands. Functional assembly of the system approximately follows second order kinetics *in vitro* and improves when cotranscribed, rather than when assembled from purified components. Split-Broccoli fluorescence is digital *in vivo* and retains functional modularity when fused to RNAs that regulate circuit function through RNA-RNA hybridization, as demonstrated with an RNA Toehold switch. Split-Broccoli represents the first functional split-aptamer system to operate *in vivo*. It offers a genetically-encoded and nondestructive platform to monitor and exploit RNA-RNA hybridization, whether as an all-RNA, stand-alone AND gate or as a tool for monitoring assembly of RNA-RNA hybrids.

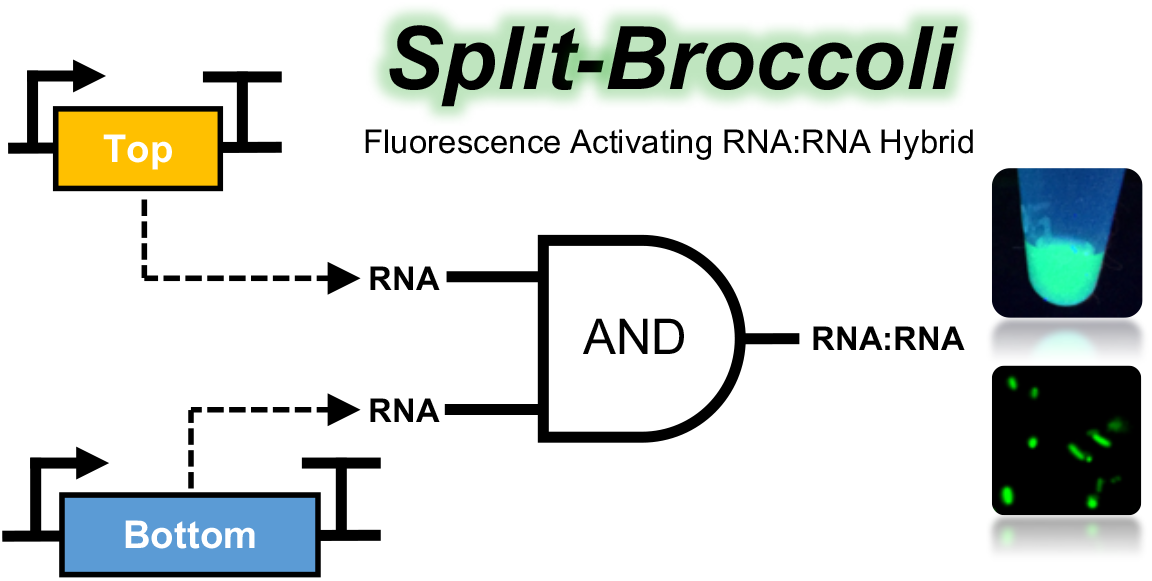

## Introduction

RNA-RNA interactions drive key processes in biology, such as RNA interference,^1^ retroviral genome dimerization,^2^ CRISPR-Cas activation^3^ and targeting,^4^ and post-transcriptional regulation of RNA by long noncoding RNAs.^5^ RNA is also increasingly utilized as a programmable element for the control of gene expression and cellular function within the field of synthetic biology.^6^,^7^ Riboregulators,^8^ Toehold switches,^9^ Small Transcription Activating RNAs (STARs),^10^ and other ribodevices exploit the hybridization of a *trans-*acting RNA with a target mRNA to control gene expression at either the transcriptional or translational level. Nevertheless, the toolkit for monitoring RNA-RNA hybridization events as they occur in real time is limited, relying primarily on fluorescent reporter proteins. However, this approach is indirect, as the downstream measurements are subject to additional variables related to translation, time delays related to fluorophore maturation, and additional resource burdens on the system. Monitoring RNA-RNA hybridization directly through an RNA-based system is therefore highly desirable. RNA-based systems have small genetic footprints and they offer improved temporal sensitivity, as signal amplification and decay would be governed by transcription, degradation and assembly rates of the RNA. Furthermore, genetic circuits with a directly reportable output driven by RNA-RNA hybridization may provide an alternative detection strategy for cell-based or cell-free diagnostics based entirely on transcription.^11^

Intracellular RNA visualization strategies roughly fall into two categories: those that are genetically encoded and those that rely on chemically-labeled probes.^12^ Genetically-encoded strategies allow for nondestructive analysis in real time, unlike chemical probes that require exogenous introduction and may thereby perturb the system. Several encoded systems for RNA visualization have been developed that utilize RNA-protein interactions, such as those found in the bacteriophages MS2 and PP7. RNA hairpin motifs in these phage genomes serve as binding sites for phage coat proteins. When these motifs are inserted into an RNA of interest they can recruit coat proteins that have been fused to fluorescent reporter proteins, allowing for indirect visualization of the RNA.^13,14^ However, these systems require the insertion of up to 24 copies (∼1200 nucleotides) of the binding motif and the use of export and localization signals to overcome diffuse fluorescence arising from unbound fusion proteins. While some of the limitations of protein-based RNA visualization approaches can be mitigated by fluorescence complementation assays—which utilize split-fluorescent proteins that only fluoresce when brought into proximity through paired RNA binding motifs—these approaches still require proteins.

Genetically-encoded RNA aptamers that bind and activate the fluorescence of small molecule dyes have seen increasing use in monitoring RNA directly without involving reporter proteins. Aptamers are single-stranded, functional nucleic acids that fold into three-dimensional structures to bind defined molecular targets.^15^ Several RNA aptamers have been selected against small molecule dyes for which fluorescence is activated upon binding and sequestration within the aptamer.^16-18^ While the malachite green aptamer (MGA)^16^ was the first to demonstrate such fluorescence activation, the recent selection of the Spinach aptamer,^17^ which binds to a freely diffusible and nontoxic dye (DFHBI), has resulted in a surge of publications reporting its use as an output for engineered genetic circuits,^19,20^ as a tool to monitor RNA transcription,^21,22^ and as a fluorescent sensor for metabolites.^23,24^ Several improvements to the original Spinach aptamer have been developed through directed evolution strategies,^25-27^ including the Broccoli aptamer, which was selected *in vivo* for improved physiological properties.^28^ The utility of these aptamers was further expanded by the introduction of DFHBI-1T, a brighter aptamer-activatable fluorophore compatible with the Spinach family of aptamers and optimized for conventional green fluorescence excitation and emission spectra used for GFP detection.^29^ Nonetheless, while several direct visualization strategies for RNA exist, including those that operate exclusively as RNA, none is designed specifically for visualization of RNA-RNA interactions.

We reasoned that RNA assembly processes could be monitored directly through the use of a split-aptamer system analogous to the split fluorescent proteins that have been used extensively to monitor cosynthesis, colocalization, and assembly of proteins. Split fluorescent aptamers could provide similar information for RNA, as they would only generate signal upon cosynthesis, colocalization, and hybridization of the component nucleic acids to form the functional aptamer. Indeed, split versions of MGA^30^ and Spinach^31-33^ aptamers have been developed for use *in vitro*. However, these systems have not been directly observed to function when transcribed *in vivo*. The triphenylmethane dye that binds to MGA is toxic, therefore limiting its use in living systems, and the Spinach aptamers are limited by weak signal *in vivo*. Multimerization can increase signal strength, but can also result in interference between the different aptamer modules. Several approaches to multimerizing or splitting functional nucleic acids utilize three-way junction (3WJ) architectures,^34-38^ which are replete throughout biology.^39^ Because the Broccoli aptamer exhibits improved physiological properties, such as lowered dependence on magnesium and higher folding efficiency at 37°C,^28^ we reasoned that it would serve as an ideal platform for an improved split-aptamer system. We further reasoned that minimal multimerization of the aptamer would overcome potential issues of low signal and that a self-assembling and thermodynamically stable RNA motif, such as a 3WJ, would allow for robust fluorescence complementation.

Here, we present the binary “Split-Broccoli” system as a stand-alone RNA logic gate and as a device for monitoring RNA-RNA hybridization *in vitro* and *in vivo*. The design-build-test cycle iterates from *in silico* design to *in vitro* implementation and finally to *in vivo* functionality. The structure-based strategy begins with an unsplit dimeric aptamer within a stabilizing RNA architecture and continues with its bisection into a two-component system. The system assembles reliably *in vitro* when purified components are thermally renatured, and the addition of transcription terminator structures improves the “OFF” level of the system. At physiological temperature, *in vitro* assembly approximately follows second-order kinetics but appears to be limited by kinetic traps that prevent fast refolding into the functional hybrid. However, when individual RNAs of the system are cotranscribed *in vitro*, fluorescence signal strength roughly approximates the unsplit variant over a 4-hr time course. The Split-Broccoli system also assembles and activates fluorescence when expressed *in vivo*, whether as a stand-alone AND gate or as a tool to monitor an RNA-RNA hybridization event that drives translation of a red fluorescent protein. We anticipate that the Split-Broccoli system is an enabling addition to the toolkit for RNA biologists.

## Results

### Designs for Stabilized Broccoli Aptamers and the Split-Broccoli System

The published monomeric and dimeric forms of the Broccoli aptamer (*mBroccoli* and *dBroccoli*, Table S1) served as starting points for engineering three dimeric forms of Broccoli aptamer.^28^ First, while the Spinach series of aptamers^17,25^ are typically embedded within a tRNA^Lys^3 scaffold^40^ to stabilize folding and *in vivo* accumulation, there are divergent reports as to whether Broccoli’s fluorescence activation requires transcription within a tRNA context.^27,28,41^ We therefore compared the performance of *mBroccoli* to *dBroccoli* without additional sequence and also to the Spinach2 aptamer embedded within a tRNA^Lys^3 scaffold (*tSpinach2*). Fluorescence from *mBroccoli* was equivalent to that of *tSpinach2,* but *dBroccoli* alone, without additional stabilizing base pairs, yielded approximately 25% of signal relative to *mBroccoli* (Figure S1). This represents an 8-fold reduction from the expected doubling, consistent with its predicted suboptimal folding (Figure S2). Alternative designs for stabilizing Broccoli dimer without greatly increasing transcript size were guided by RNA secondary structure predictions using the web-based NUPACK software suite.^42^ Adding 3 terminal G:C base pairs to the terminal stem—but not 1 or 2 G:C base pairs or 3 pairs that included A:U pairs—provided adequate stabilization to favor the expected structure, but stabilization by a mixed 4 base pair stem (GAGG:CTCC) exceeded that of the other designs. When this 4 base pair stem was appended to *dBroccoli* to form the 100-nucleotide “stabilized dimeric Broccoli” (*SdB*) (Figure S2d), fluorescence was more than 2-fold enhanced relative to *mBroccoli* (Figure S3). *SdB* was therefore used as a reference point for further engineering of the aptamers.

The second dimeric Broccoli aptamer design exploits the 3WJ motif derived from the packaging RNA (pRNA) component of the DNA packaging motor from bacteriophage Φ29. This stable element is a proven architecture for multimerization of functional nucleic acids^35,43,44^ and self-assembly from oligonucleotides.^45-47^ We therefore inserted Broccoli monomers into arms 2 and 3 of the 3WJ to form *3WJdB* (Figure 1a). Because these arms extend in opposite directions from the 3WJ structural core in crystal structures of the Φ29 3WJ pRNA,^48^ this design is expected to keep the monomer units spatially distant and to minimize intersubunit misfolding or potential quenching that can arise from chromophore-chromophore interactions. While the work reported here was under way, Filonov *et al*. reported a similar design (“*F30-2xBroccoli*”) utilizing the same 3WJ element from Φ29 pRNA, but with Broccoli monomers inserted into arms 1 and 2.^49^ Our *3WJdB* design incorporates a single nucleotide substitution (C → U) and conversion of a U:A base pair to an A:U pair, as Filonov *et al.* observed that these changes improved transcriptional yield of full-length RNA by interrupting a poly-uridine tract that may serve as a cryptic transcription terminator.^49^ However, we also note that these substitutions include an adenosine whose phosphate is implicated in coordinating one of the core metal ions of the 3WJ,^48^ potentially affecting thermostability of the motif.

The third dimeric Broccoli aptamer design bisects *3WJdB* after first inverting the monomer in Arm 2 to ensure that neither strand alone would contain the full sequence required to form a functional aptamer. In addition, 8 or 12 base pairs were appended on the two distal stems to favor complete hybridization of the two strands and formation of the desired secondary structure (Figure 1b). The two autonomous RNA strands of the Split-Broccoli system are designated *Top* and *Bottom*. Neither strand contains the full sequence required to form a functional Broccoli monomer, and neither is predicted to fold into a secondary structure that resembles a functional monomer (Figure S4a, b). Engineering the Split-Broccoli system in this manner, wherein fluorescence activation (output) is only expected in the presence of both *Top* and *Bottom* strands (inputs), creates an AND gate capable of performing and reporting on logical operations entirely as RNA (Figure 1c).

**Figure 1.**
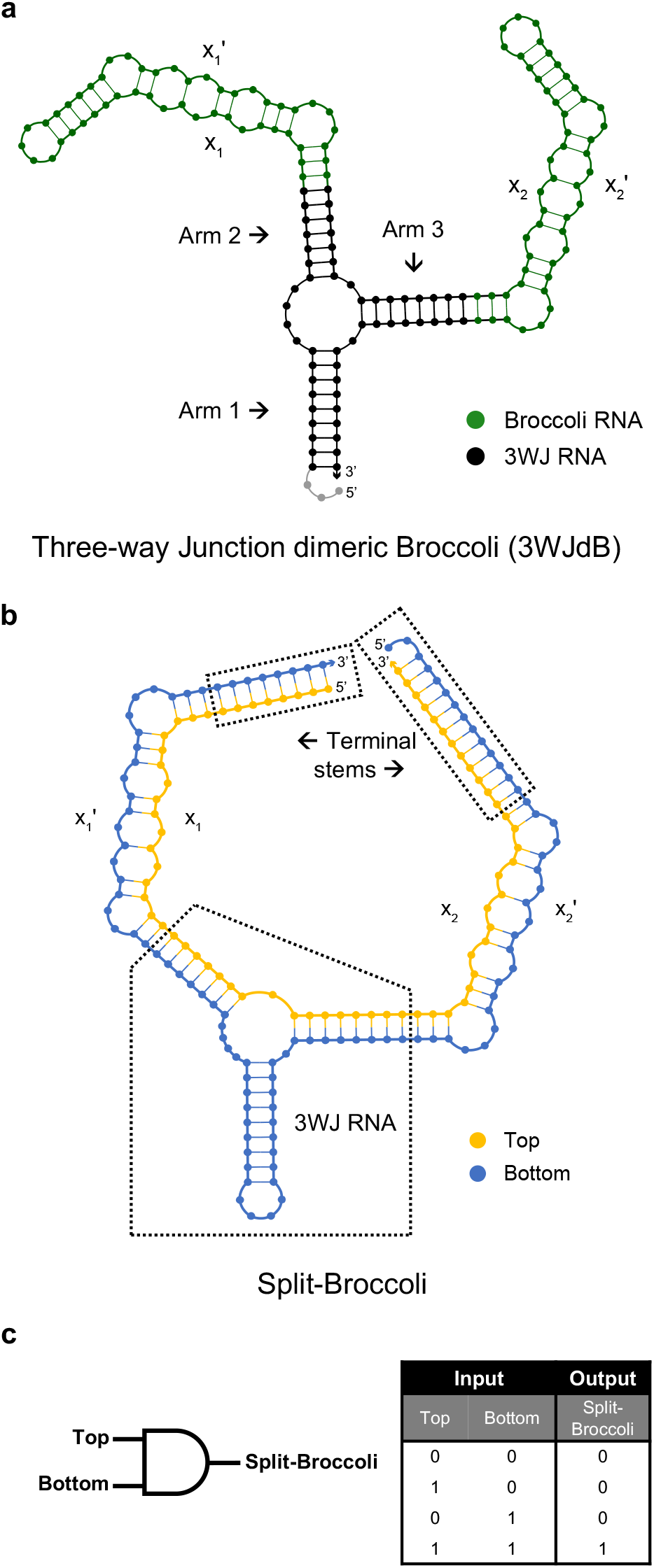
Design and NUPACK predicted secondary structures of *3WJdB* and *Split-Broccoli*. (**a**) Monomers of Broccoli aptamer (green) were inserted into Arms 2 and 3 of the three-way junction (3WJ) RNA motif (black) to create the unimolecular, unsplit three-way junction dimeric Broccoli (*3WJdB*). (**b**) Design of the Split-Broccoli system required inversion of the Broccoli monomer present in Arm 2 to ensure that neither *Top* (yellow) nor *Bottom* (blue) strand alone contained the full sequence required to independently form a fully functional monomer of Broccoli (i.e. x_n_:x_n_’). Predicted hybridization of the two strands was strengthened with the addition of terminal stems. (**c**) The Split-Broccoli system illustrated as an RNA AND gate (left) and its corresponding truth table (right). Output from the system should be true (1) only when both inputs are also true.

### *Split-Broccoli* Assembles *In Vitro* to Reconstitute Fluorescence Activation

To evaluate performance of the Split-Broccoli system, we compared two approaches for assembling *Split-Broccoli* from purified *Top* and *Bottom* strands. In the first approach, equimolar amounts of *Top* and *Bottom* were mixed and incubated with dye in buffer, then unfolded at 90°C and renatured by slow cooling to 37°C prior to measuring fluorescence. The thermally renatured *Split-Broccoli* exhibited approximately 82% of the signal relative to the unsplit, unimolecular *3WJdB*, while neither *Top* nor *Bottom* strand alone generated robust fluorescence signal (Figure 2a). Fluorescence signal for the renatured *Split-Broccoli* was 22-fold greater than *Bottom* strand alone, which gave less than 4% of signal compared to *3WJdB*. Additionally, the *3WJdB* design exhibited higher signal when compared to *SdB*, suggesting that the 3WJ framework reinforces the productive fold better than the minimal stem that stabilizes *SdB* (Figure 2a). In the second approach, the two strands were folded separately and mixed at 37°C without renaturation to mimic a more biologically relevant assembly process. Fluorescence signal for the complex was again strongly above background, and little or no fluorescence was observed for either *Top* or *Bottom* strand alone (Figure 2b). These patterns were clearly visible when these samples were excited with a standard UV-light source (Figure 2c). For this simple mixing approach, fluorescence from the assembled complex was approximately 60% of the signal from unsplit *3WJdB* under the same conditions and was roughly 15-fold over *Bottom* strand alone, which again generated ∼4% signal over the *No RNA* background sample (Figure 2b). The data from these two approaches show effective reconstitution of aptamer function from the binary Split-Broccoli system *in vitro*, with switch-like behavior that serves as a two-input AND gate and fluorescence signal strength that is slightly reduced relative to the unsplit *3WJdB* and roughly equivalent to *SdB*.

**Figure 2.**
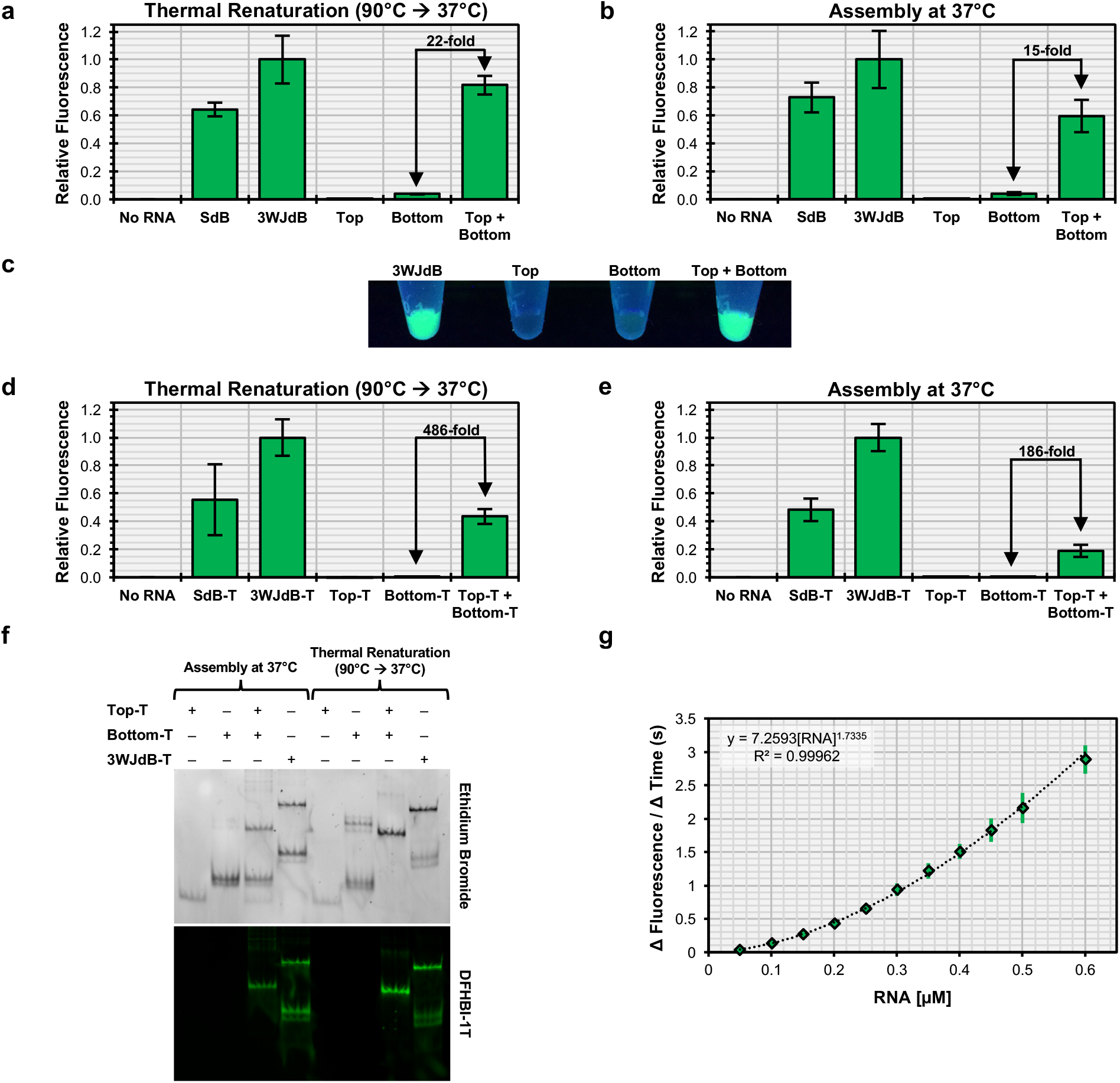
*In vitro* assembly of the Split-Broccoli system in the absence and presence of transcription terminator structures. Assembly of equimolar amounts of purified Split-Broccoli RNA components (*Top* + *Bottom*) demonstrate robust function comparable to fluorescence of the stabilized dimeric Broccoli (*SdB*) and *3WJdB*, (**a**) when thermally renatured or (**b**) when simply incubated together at physiological temperature. Background signal from either *Top* or *Bottom* alone remains minimal for both assembly methods. (**c**) Fluorescence of *3WJdB* and the Split-System (*Top* + *Bottom*, thermally renatured) is easily observed when excited with longwave ultraviolet light, whereas signal from *Bottom* alone is only slightly discernable. When transcribed with transcription terminator structures (denoted by appending *“-T*” to the names of the individual RNAs) and assembled *in vitro* (**d**, **e**), the Split-Broccoli system exhibits a decrease in relative fluorescence, but demonstrates a larger fold-change in fluorescence activation over either *Top-T* or *Bottom-T* alone. (**f**) Non-denaturing gel electrophoresis and dual staining with ethidium bromide and DFHBI-1T of the Split-Broccoli system with transcription terminator structures suggests that decreased fluorescence of the system is a result of incomplete hybridization between *Top-T* and *Bottom-T*, rather than nonfunctional assembly. (**g**) Functional assembly of *Top-T* and *Bottom-T* approximately follows second-order kinetics (y = A[Top][Bottom] = A[Top]^2^, for equimolar mixture). Mean values are shown with error bars to indicate standard deviation (n = 5 for panels **a, b, d, e**; n = 4 for panel **g**).

### Transcription Terminator Structures Enhance Activation Ratio

The RNA species described above were generated by run-off transcription of linear dsDNA templates. In contrast, bacterial Rho-independent transcription termination utilizes a G:C rich, stem-loop structure followed by a U-rich tract that promotes dissociation of the polymerase from the DNA template.^50^ To characterize the performance of Split-Broccoli in a more biological sequence context, T7 transcription terminator structures were appended to both unsplit and split aptamer transcripts, denoted by appending *“-T*” to the names of the individual RNAs. Neither *Top-T* nor *Bottom-T* was predicted to contain a Broccoli-like secondary structure (Figure S4c,d), whereas hybridization of these two strands was predicted to form the two Broccoli domains flanking the 3WJ motif (Figure S5). Fluorescence from thermally renatured *Top-T* and *Bottom-T* strands was 486-fold above signal for either strand alone, representing switch-like, digital behavior for the Split-Broccoli system. In comparison with runoff-transcripts, this improvement in activation ratio can be ascribed to the reduction in signal from *Bottom-T* alone to background *No RNA* levels, in spite of a decrease in signal from the assembled complex relative to the unsplit *3WJdB-T* (∼43%, Figure 2d). Fluorescence activation remained high (186-fold) when the complex was assembled by incubating *Top-T* and *Bottom-T* strands at physiological temperature, even though signal strength for the complex dropped to approximately 19% of the unimolecular *3WJdB-T* (Figure 2e). Fluorescence signal strength for unsplit *3WJdB and 3WJdB-T* were roughly equivalent under both conditions (Figure S6), thus justifying comparison of relative fluorescence values of terminated transcripts to run-off transcripts.

### Split-Broccoli Assembly Approximately Follows Second Order Kinetics

To address the molecular basis for limitations to the *in vitro* assembly of *Top-T* and *Bottom-T* strands at physiological temperature, assembly of *Split-Broccoli* was evaluated using a native gel stained with ethidium bromide, to locate all molecular species, and DFHBI-1T, to identify functionally assembled complexes.^49^ When incubated at 37°C, each individual strand of the Split-Broccoli system migrated as a single nonfunctional band, and a new functional band appeared when the two strands were assembled together (Figure 2f, **S7**). However, a substantial amount of input material remains unassembled under these conditions, consistent with the observed decrease in relative fluorescence for the annealed complex upon adding transcription terminator structures (Figure 2e). Annealing efficiency and fluorescence intensity of the system improved when the system was thermally renatured. Although each strand remained independently nonfunctional when renatured, a second isoform of *Bottom-T* became prominent, indicating that the terminator hairpins may be cross-annealing to form a dimer of *Bottom-T*. Together, these results suggest that incomplete, rather than nonfunctional, assembly is responsible for the decrease in signal when purified components of the system are incubated at 37°C. Robust assembly *in vitro* may thus be limited by kinetic folding traps within one or both strands that prevent fast refolding into the functional hybrid.

In principle, RNA-based genetic control elements can provide rapid responses *in vitro* or *in vivo* because signal development does not require translation. To better understand assembly kinetics, we measured the rate of functional assembly of Split-Broccoli at physiological temperature. *Top-T* and *Bottom-T* strands were separately folded, mixed with dye, and mixed with each other at equimolar concentrations. Development of fluorescence signal was monitored as a function of time (Figure S8), and the rates were plotted against input RNA concentration. *Split-Broccoli* assembly approximately followed second order kinetics (Figure 2g). The exponent (1.73) falls slightly below the expected value of 2.00, potentially as a consequence of competition from internal structure within the individual strands, as suggested by the native gel mobility shift assay described above.

### Cotranscriptional Assembly of Split-Broccoli Improves Signal

Gel purification of RNA under non-native conditions can introduce conformational traps.^51^ In contrast, cotranscriptional folding *in vivo* is a sequential process in which the RNA folds from the 5’ end as synthesis occurs. Furthermore, folding is concurrent with, and can be influenced by, other RNA species that are present. Therefore, to better characterize *in vitro* the *in vivo* potential of the Split-Broccoli system, we evaluated cotranscriptional assembly by incubating equimolar amounts of linear dsDNA templates corresponding to *3WJdB-T*, *Top-T*, *Bottom-T*, or both *Top-T* and *Bottom-T* together, and assaying for function in transcription reaction conditions optimized for time resolution (Figure 3). The unimolecular *3WJdB-T* exhibited measurable fluorescence within a few minutes of incubation, and *Split-Broccoli* required approximately ten more minutes as the two strands accumulated to a sufficient concentration to associate with each other at a detectable level. Although the rate of signal development for the unimolecular *3WJdB-T* is substantially faster than *Split-Broccoli* for the first forty-five minutes, the rates are approximately equal for both the unsplit and split aptamers afterwards. Additionally, when compared to *SdB-T* in transcription conditions optimized for RNA production, *3WJdB-T* function is slightly improved and goes to completion, suggesting that the inclusion of a 3WJ scaffold does not delay aptamer folding (Figure S9). Cotranscription of the split-aptamer system reconstituted nearly all of the function of *3WJdB* (∼88%) and demonstrated a remarkable 124-fold increase in signal relative to *Bottom-T* strand alone after a 4-hr reaction. Cotranscription of the Split-Broccoli system with terminator structures therefore allows for robust functional assembly without the requirement for additional *in vitro* manipulations, such as thermal renaturation or coincubation of independently transcribed RNA that may be kinetically trapped in alternative, nonfunctional structures.

**Figure 3.**
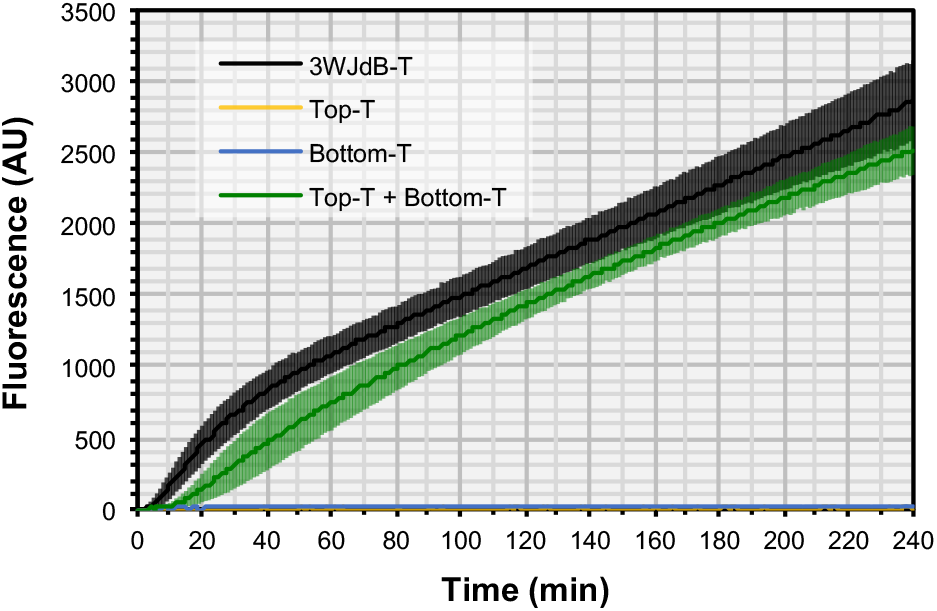
Cotranscriptional assembly of *Split-Broccoli* with transcription terminator structures improves signal relative to independently transcribed and assembled RNA. When cotranscribed with equimolar amounts of template DNA in a one-pot *in vitro* transcription reaction designed to maximize time resolution, *Split-Broccoli* (*Top-T + Bottom-T*, green) exhibits approximately 88% of the signal generated by the unimolecular, unsplit variant (*3WJdB-T*, black) after a 4-hr reaction. By the final time point at 240 min, Split-Broccoli demonstrates a 124-fold increase over either *Top-T* (yellow) or *Bottom-T* (blue) alone. Mean values are shown (n = 4) with error bars to indicate standard deviation.

### Split-Broccoli Functions *In Vivo*

To determine whether the Split-Broccoli system would function *in vivo*, we constructed expression plasmids for evaluation in *Escherichia coli*. Transcripts originate from the constitutive P70a promoter^52^ and were terminated with Rho-independent transcription terminators. Single-insert plasmids direct the transcription of either *3WJdB-T* (pUC19-P70a-3WJdB-T), *Top-T* (pUC19-P70a-Top-T) or *Bottom-T* (pUC19-P70a-Bottom-T), while a dual-insert plasmid (pUC19-P70a-Top-T∼P70a-Bottom-T) directs transcription of *Top-T* and *Bottom-T*, with a 270-nucleotide spacer (∼) between transcription units to reduce topological tension arising from simultaneous transcription (Figure 4a and **Supporting Information**). As a control, we also included a variant of this plasmid that was missing the second P70a promoter upstream of *Bottom-T* (pUC19-P70a-Top-T∼Bottom-T) to ensure that signal generated by the dual-promoted Split-Broccoli system is not a product of a single run-on transcript folding onto itself.

Transformed bacteria were grown to mid-log phase and their fluorescence was measured by flow cytometry (Figure 4b,c). For the cell population containing the *3WJdB-T* insert, mean fluorescence intensity was shifted approximately 25-fold (Figure 4b) relative to pUC19 transformants alone. Importantly, transcribing both *Top-T* and *Bottom-T* together generated a notable shift (approximately 6-fold above background) in mean fluorescence intensity. Fluorescence signal from populations containing only *Top-T* or *Bottom-T* inserts exhibited background levels of signal equivalent to the unmodified pUC19 plasmid containing no insert, representing a true OFF state, as did populations transformed with a plasmid that lacked the second promoter for transcribing *Bottom-T,* indicating that transcription of *Top-T* strand was effectively terminated (Figure 4b,c). These results were further confirmed with fluorescence microscopy (Figure 4d). The Split-Broccoli system therefore acts as a stand-alone, all-RNA AND gate *in vivo*, with a true OFF state in the absence of both strands and an ON state in their presence.

**Figure 4.**
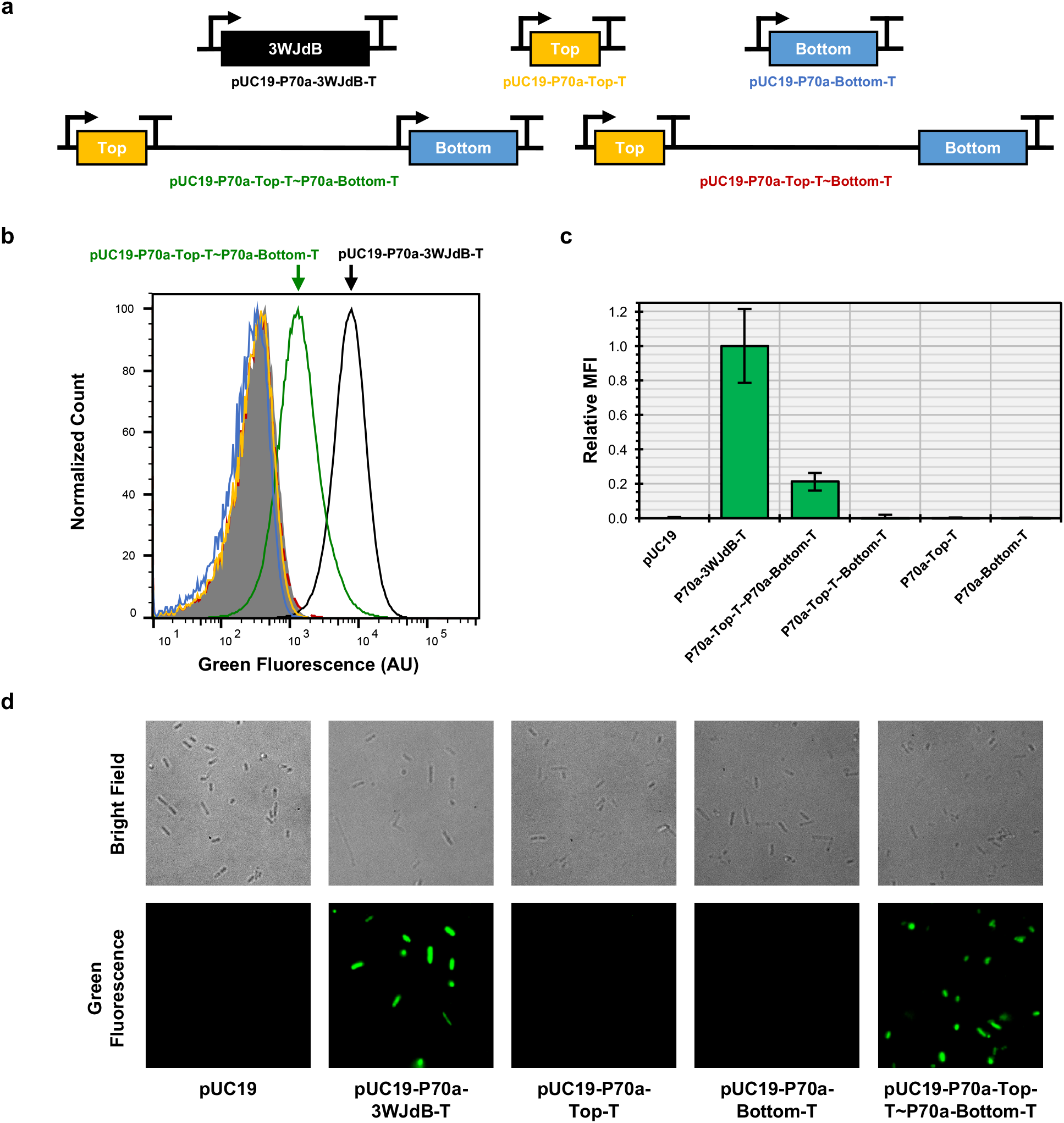
The Split-Broccoli system functions when expressed *in vivo*. (**a**) DNA templates corresponding to *3WJdB-T* (black), *Top-T* (yellow), *Bottom-T* (blue) were individually cloned into the pUC19 plasmid. A single plasmid expressing both *Top-T* and *Bottom-T* was created (pUC19-P70a-Top-T∼P70a-Bottom-T), as was a control plasmid for run-on transcription which lacked a promoter immediately upstream of *Bottom-T* (pUC19-P70a-Top-T∼Bottom-T). (**b**) A representative flow cytometry histogram of 5 x 10^4^ events per population illustrates a shift in fluorescence for the plasmid containing the Split-Broccoli expression plasmid (green). Bacterial populations transformed with plasmids containing either *Top-T* or *Bottom-T* alone, or lacking a promoter upstream of *Bottom-T*, demonstrate background levels of fluorescence equivalent to the unmodified pUC19 plasmid control. (**c**) Relative mean fluorescence intensities for flow cytometric analyses of transformed populations, normalized to the pUC19 plasmid (0) and *3WJdB-T* (1), are shown with error bars to indicate standard deviations (n ≥ 4). (**d**) Fluorescence microscopy imaging further validates the *in vivo* functionality of the Split-Broccoli system, as green fluorescence is only observed for *E. coli* transformed with either the unimolecular *3WJdB-T* encoding plasmid or bimolecular *Split-Broccoli* encoding plasmid.

### Split-Broccoli Is Functionally Modular and Can Be Used to Monitor RNA-RNA Assembly *In Vivo*

RNA molecules assemble with each other to drive many processes in biology. To evaluate whether the Split-Broccoli system could report on such assembly events, we chose to monitor activation of an RNA Toehold switch. Toehold switches are two-component RNA systems for gene regulation in which a *trans*-acting RNA (“*Trigger*”) relieves translational repression from a *cis*-acting hairpin structure (“*Toehold*”) that lies just upstream of a gene of interest.^9^ In the absence of a *Trigger* RNA, the *Toehold* structure effectively sequesters ribosomal access to the 5’ UTR. Base pairing of *Trigger* to *Toehold* relaxes the 5’ UTR hairpin structure, thereby exposing the ribosome binding site and start codon to allow for efficient translation initiation. We fused *Top* with a well-characterized *Toehold* that regulates an mCherry reporter protein to generate *Top-Toehold-mCherry* and we fused *Bottom* with the corresponding *Trigger* RNA to generate *Trigger-Bottom* (Figure 5a,b), and then determined fluorescence for bacteria carrying these constitutively-expressing plasmids. Red fluorescence for bacteria encoding *mCherry* in the OFF state (*Top-Toehold-mCherry* only) showed minimal red fluorescence, potentially due to incomplete suppression of translation by the Toehold (Figure 5c, S10). Both *Top-Toehold-mCherry* alone and *Trigger-Bottom* alone exhibited less than 2% of green fluorescence of the functional *Top+Bottom* populations. Finally, when the complete Split-Broccoli system was fused with the complete Toehold switch system and expressed as two discrete transcripts (*Top-Toehold-mCherry* + *Trigger*-*Bottom*), both green and red fluorescence were activated in the cell population and observed with flow cytometry (Figure 5c, S10) and fluorescence microscopy (Figure 5d). Split-Broccoli is therefore highly modular and can be used to monitor RNA-RNA interactions *in vivo*.

**Figure 5.**
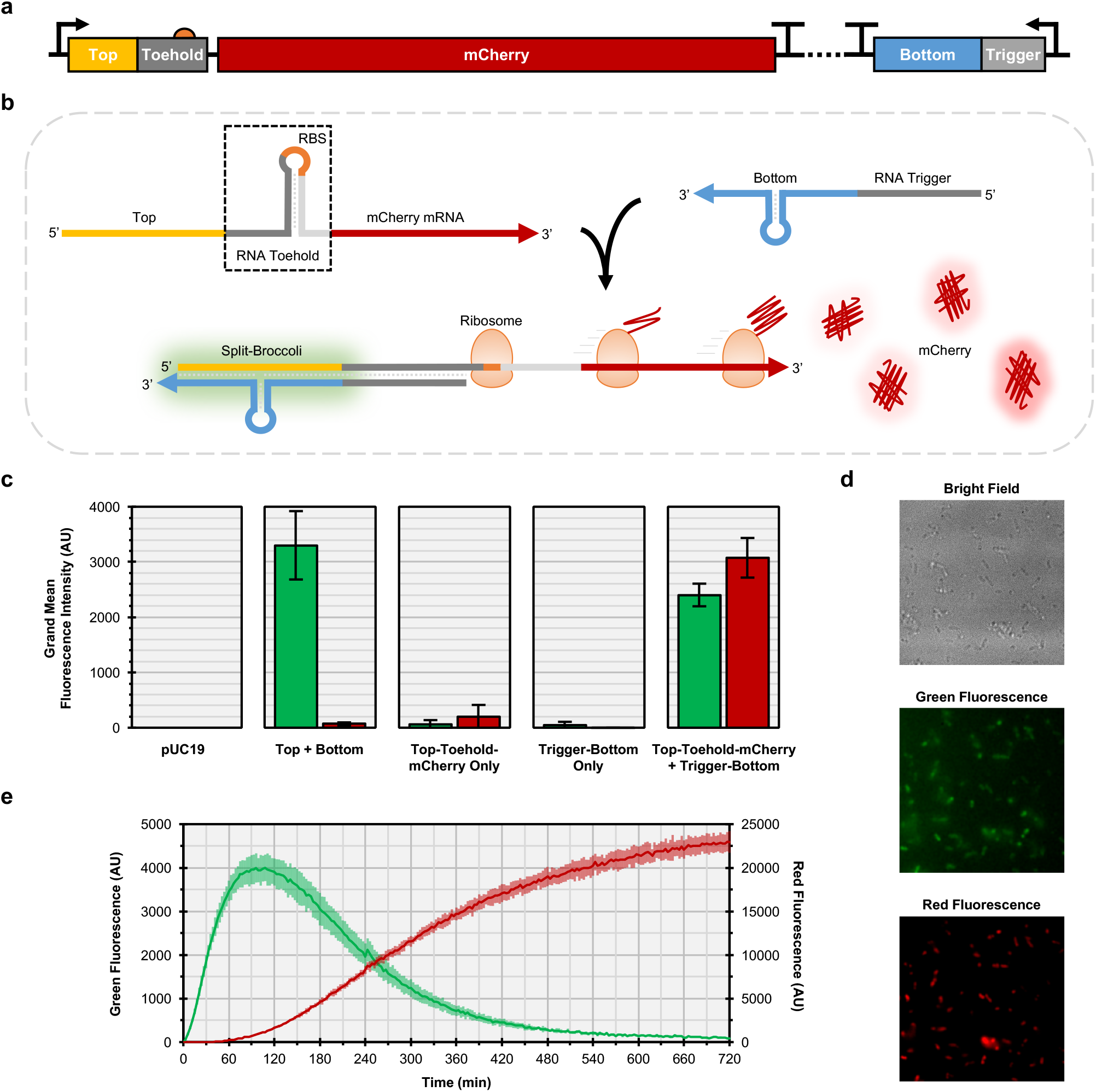
Split-Broccoli is modular and can be used to monitor RNA-RNA hybridization events *in vivo*. (**a**) A Split-Broccoli Toehold Switch plasmid was constructed to include two constitutively-expressed transcription units. The first transcription unit encodes *Top* (yellow) and *Toehold* (gray) sequences within the 5’ UTR of the *mCherry* mRNA (red). Translation of the *Top-Toehold-mCherry* mRNA is suppressed due to sequestration of the ribosome binding site (orange) and start codon within the toehold structure (boxed). The second transcription unit encodes *Trigger* (gray) and *Bottom* (blue) sequences, which can base pair with *Top-Toehold-mCherry*. (**b**) Hybridization of *Top-Toehold-mCherry* with *Trigger-Bottom* allows fluorescence activation of the Split-Broccoli system and translation of *mCherry*. (**c**) Green fluorescence (left columns) and red fluorescence (right columns) from flow cytometric analysis of populations show background levels of fluorescence for plasmids encoding a single transcription unit only (*Top-Toehold-mCherry* or *Trigger-Bottom*). *Top + Bottom*, which transcribes the Split-Broccoli system, exhibits only green fluorescence, while the Split-Broccoli Toehold Switch plasmid (*Top-Toehold-mCherry* + *Trigger-Bottom*) exhibits both red and green fluorescence, indicating both hybridization of Split-Broccoli and translation of *mCherry*. Grand mean fluorescence intensity (n = 4) is shown with error bars to indicate standard deviation. (**d**) Fluorescence microscopy imaging of *E. coli* harboring the Split-Broccoli Toehold Switch plasmid confirms hybridization of the Top and Bottom components of *Split-Broccoli* (green fluorescence) and activation of *mCherry* translation (red fluorescence). (**e**) An *E. coli* cell-free system (TX-TL) was used to monitor transcription and translation of the Split-Broccoli Toehold Switch plasmid and demonstrates the increased temporal sensitivity of *Split-Broccoli* (green fluorescence, left axis) over *mCherry* (red fluorescence, right axis). Mean values (n = 3) are shown with error bars to indicate standard deviation.

An advantage of RNA-based fluorescent reporters over proteins such as GFP and mCherry is improved temporal sensitivity, as translation and fluorophore maturation would not be required for function. To investigate how quickly *Split-Broccoli* could be observed relative to a translated gene of interest, we utilized an *E. coli* cell-free transcription and translation system (TX-TL)^52^ with the Split-Broccoli Toehold Switch plasmid (*Top-Toehold-mCherry* + *Trigger*-*Bottom*). The TX-TL system is prepared from a crude *E. coli* cytoplasmic extract and contains the endogenous transcription and translation machinery, allowing for rapid *in vitro* characterization of synthetic gene circuits with *in vivo*-like conditions. Green fluorescence from *Split-Broccoli* was detectable in TX-TL almost immediately and increased rapidly before peaking around 90 minutes (Figure 5e). In contrast, red fluorescence from *mCherry* took approximately 45 minutes to detect and slowly increased over the remaining time course. While the fluorophore maturation time can be improved through the use of alternative fluorescent proteins, the nearly instantaneous response of the Split-Broccoli system demonstrates the speed at which an RNA-based reporter can act.

In this application of the Split-Broccoli system, the RNA-RNA hybridization event self-reports through green fluorescence of the assembled aptamer, while translational activation of the downstream *mCherry* gene is subsequently self-reported through red fluorescence. Most proteins do not self-report their synthesis and are not so easily monitored in real time. The *Top-Toehold-mCherry* and *Trigger-Bottom* design can be readily adapted to those proteins by replacing *mCherry* with another gene of interest for which translation is less readily monitored. The appearance of green fluorescence upon induction of *Trigger-Bottom* could then be taken as visual evidence that the gene of interest was being translated. Although we fused *Split-Broccoli* to an RNA Toehold switch, alternative systems for regulating gene expression could also be used, such as those regulated by Small Transcription Activating RNAs (STARs).^10^ The hybridization of an antisense STAR to its cognate sense RNA results in the formation of an anti-terminator structure that allows transcription of an mRNA to continue. The Split-Broccoli system could therefore be used to monitor the STAR hybridization event upstream of a gene of interest.

## Discussion

The 3WJ RNA architecture is a robust platform for dimerization of the Broccoli aptamer and allows for a straightforward bisection of aptamers into two-stranded systems that can restore function when hybridized. We have shown that the Split-Broccoli system functions when assembled from purified RNA *in vitro*, in both the presence and absence of transcription terminator structures and when thermally renatured or assembled at physiological temperature. *In vitro* performance appeared to be limited by an incomplete assembly process that was only partially restored through thermal renaturation, suggesting a high kinetic barrier for refolding of the RNA components into the hybridized complex. This barrier was inconsequential if the components of the system were cotranscribed, allowing for the more native RNA structures to hybridize efficiently and productively. These findings motivated us to explore the use of the system *in vivo*. Although Split-Broccoli signal *in vivo* was observable, it was not as robust as the unimolecular *3WJdB* construct. We speculate that this reduction in signal could have arisen from instability of one or both of the strands against degradative cellular forces, such as RNases, rather than kinetic barriers to refolding. The decrease in relative signal could potentially be remediated by transcribing the split aptamer system with highly structured 5’ ends, as the 3’ end terminator structures likely offer some protection against nuclease activity targeting unstructured, linear RNA. Furthermore, when the Split-Broccoli system was fused to an RNA Toehold switch upstream of a fluorescent protein, *Split-Broccoli* fluorescence was observed tens of minutes before the fluorescent protein, demonstrating the speed and utility of an RNA-based split fluorescent aptamer in monitoring an RNA-RNA hybridization event.

To the best of our knowledge, Split-Broccoli is the first split-aptamer system directly shown to function when transcribed *in vivo* and is the first example of an *in vivo* logic gate with RNA inputs and a directly observable output composed of RNA. We posit that Split-Broccoli, or an analogously composed system, can be extended to detect RNA-RNA hybridization events nondestructively in real time as an alternative to chemically synthesized probes and without the requirement of a protein reporter. For RNA-based computation strategies that require a measureable reporter in living systems, Split-Broccoli may offer an approach that can operate without introducing an additional time delay or resource burden required by translating traditional protein-based reporters. Additionally, as transcription of the individual strands of the Split-Broccoli system can be regulated by independent transcription factors, the Split-Broccoli system can be used as an AND gate *in vivo*, further expanding the functional capabilities of RNA as a programmable and self-reporting molecule for RNA-RNA assembly.

## Methods

### Broccoli Designs and NUPACK Analysis

Sequences for the Spinach2 aptamer, Broccoli aptamer and the core domain of the Φ29 pRNA 3WJ motif were used as previously published.^25,28,45^ *SdB*, *3WJdB* and *Split-Broccoli* designs, with and without transcription terminators, were analyzed for predicted structures with the NUPACK web application at default settings for RNA at 37°C.^42^ For Split-Broccoli hybridization, the two input strands were constrained to a maximum complex size of 2 and at a concentration of 1 μM each. For details of DNA sequences refer to Supporting Information.

### DNA Templates and RNA Transcription

Oligonucleotides for generating DNA templates were ordered from Integrated DNA Technologies and ligated to generate PCR amplification templates. In short, oligonucleotides corresponding to the left and right halves of each sequence, and a reverse complement sequence bridging both halves, were designed. The oligonucleotides corresponding to the right half of each sequence were phosphorylated for ligation using T4 polynucleotide kinase (New England Biolabs). All three oligonucleotides per complete sequence were then incubated in equimolar amounts and annealed through a heat-cool step prior to ligation with T4 DNA ligase (New England Biolabs). Ligated oligonucleotides were PCR amplified with Pfu DNA polymerase, a forward primer to add the 5’ XbaI restriction site and T7 promoter, and a reverse primer to append the T7 terminator and 3’ XmaI restriction site. Amplification products were cloned into the pUC19 plasmid and inserts were confirmed through sequencing. Plasmids encoding the T7 RNA polymerase promoted templates were deposited to Addgene (IDs 87307-87310). For details of DNA sequences refer to Supporting Information.

Transcription templates were generated by PCR amplification of sequence-verified plasmids. The pET28c-Spinach2 plasmid (a gift from Samie Jaffrey) was used for PCR amplification and transcription of *tSpinach2*. Run-off transcription reactions were performed using T7 RNA polymerase, *in vitro* transcription buffer (50 mM Tris-HCl pH 7.5, 15 mM MgCl_2_, 5 mM DTT, and 2 mM spermidine), and 2 mM of each ATP, CTP, GTP and UTP. Reactions were incubated at 37°C for a minimum of 4 hr and halted with the addition of 2X RNA loading buffer (95% formamide and 5 mM EDTA, with trace amounts of Xylene Cyanol FF and Bromophenol Blue). RNAs were purified through denaturing polyacrylamide gel electrophoresis (0.75 mm 6% TBE-PAGE, 8 M urea) and bands corresponding to the expected product size were gel extracted and eluted while tumbling overnight in 300 mM Sodium Acetate pH 5.4. Eluates were ethanol precipitated, resuspended in buffer (10 mM Tris-HCl pH 8.0, 1 mM EDTA), and stored at -20°C until further use. RNA concentrations were determined on a NanoDrop 1000 spectrophotometer (Thermo Scientific).

### *In Vitro* Assembly and Fluorescence Assays

*In vitro* assembly reactions were prepared on ice with 25 picomoles of each RNA and 250 picomoles of DFHBI-1T dye (Lucerna Technologies) in buffer (40mM HEPES pH 7.5, 100mM KCl, 1mM MgCl_2_) at a final volume of 50 μL. For thermal renaturation, samples were transferred into a preheated aluminum insert within a dry heat block set to 90°C. Following addition of samples, the aluminum insert was immediately removed from the block heater and placed on the lab bench to cool to 37°C before measurement. Assembly at 37°C was performed by placing samples in a dry heat block at 37°C for 15 min before measurement. Samples were then transferred into a clear, flat-bottom 96-well plate and measured for fluorescence (λ_ex_ = 472 nm, λ_em_ = 507 nm) on an EnSpire Mulitmode plate reader (PerkinElmer). Measurements from *No RNA* samples, which contained buffer and dye, were averaged to establish background signal. Background signal was subtracted from all measurements and individual readings were normalized for fluorescence relative to 3WJdB. Normalized values were used to calculate mean and standard deviation.

### Dual Staining Native Gel Shift Assay

10 picomoles of each purified RNA (*Top-T*, *Bottom-T*, *3WJdB-T*) were incubated at 37°C for 15 min before being loaded onto a 0.75 mm 6% native TBE polyacrylamide gel in a final volume of 25 μL with 20% glycerol. Approximately 1 hr after electrophoresis at 20 W in 4°C, the gel was stained according to an in-gel imaging protocol.^49^ In brief, the gel was stained with 5 μM of DFHBI-1T at RT for 15 min and then imaged using a Typhoon FLA 9000 (GE Healthcare) with Alexa Fluor 488 settings (473 nm laser excitation, Y520 emission filter). A destaining step was then performed on the gel with 2 washes in ultrapure Milli-Q water (EMD Millipore) for 5 min each, followed by a 5 min incubation in ethidium bromide at 0.5 μg/mL. The gel was then reimaged on a Typhoon FLA 9000 (GE Healthcare) using the ethidum bromide setting (532 nm laser excitation, O580 emission filter). Densitometry analysis was performed in Fiji.^53^ Following a slight linear contrast adjustment, the intensity value of each band was estimated. The hybridization yield for the annealed complex was calculated according to the following formula: [(annealed complex)/(free *Top-T* + free *Bottom-T* + annealed complex)].

### Split-Broccoli Assembly Kinetics

Purified samples of *Top-T* and *Bottom-T* RNA at various concentrations (0, 0.05, 0.1, 0.15, 0.2, 0.25, 0.3, 0.35, 0.4, 0.45, 0.5 and 0.6 μM) were independently prepared on ice, in buffer (40mM HEPES pH 7.5, 100mM KCl, 1mM MgCl_2_) containing DFHBI-1T (Lucerna Technologies) at 4 μM. Samples were aliquoted into a clear, flat-bottom 96-well plate and preincubated at 37°C for 15 min. Assembly was initiated by using a multi-channel pipette to transfer 50 μL of *Top-T* samples into 50 μL of *Bottom-T* samples at equimolar concentrations. Fluorescence measurements (λ_ex_ = 485/25 nm, λ_em_ = 535/25 nm) were taken on an Infinite F200 Pro plate reader (Tecan) every 10 sec for 3600 sec. The linear region between 500 and 1000 sec was used to fit a line and determine the slope (i.e. change in fluorescence as a function of time). Slope values from replicate experiments were averaged and then plotted against their concentration to obtain a value for the rate of assembly.

### Cotranscription Fluorescence Assay

Cotranscription assays were performed with PCR amplification products purified through agarose gel electrophoresis (GeneJET Gel Extraction Kit, Thermo Scientific) to isolate bands corresponding to the desired DNA template size. Templates were quantified on a NanoDrop 1000 spectrophotometer (Thermo Scientific). To ensure equivalent transcriptional burden on the RNA polymerase across samples, each sample contained a total of 10 picomoles of T7-promoted template DNA. For 3WJdB, *Top-T* alone and *Bottom-T* alone, 5 picomoles of each DNA template was added to 5 picomoles of DNA template for an unrelated RNA aptamer (80N 433min2) with no observed fluorescence activation of DFHBI-1T. Split-Broccoli sample was composed of 5 picomoles of each *Top-T* and *Bottom-T* DNA templates, while background sample contained 10 picomoles of the 80N 433min2 control aptamer template. Samples were incubated with 2.5 nanomoles of each nucleotide triphosphate (ATP, CTP, GTP, UTP), 1 unit of inorganic pyrophosphatase (New England Biolabs), 1 nanomole of DFHBI-1T (Lucerna Technologies), and 40 units of T7 RNA polymerase with its supplied buffer (New England Biolabs) in a total volume of 50 μL. Samples were prepared and kept on ice until transfer to a flat-bottom 384-well plate and measurement on a Synergy HT plate reader (BioTek) pre-warmed to 37°C. Plate reader was configured to take readings at 1 minute intervals over a period of 4 hr (λ_ex_ = 485/20 nm, λ_em_ = 528/20 nm). For supplemental information Figure S9, samples volumes were reduced to 20 μL and T7 polymerase was used at 100 U, while all other amounts remained the same. Background measurement values at each time point were subtracted from sample values before calculating mean and standard deviation.

### *In Vivo* Fluorescence

Expression plasmids for *in vivo* assays were constructed to contain constitutive *E. coli* P70a promoters^52^ and either T7 transcription terminator or a derivative of the T500 transcription terminator from bacteriophage Φ82.^54^ Plasmids for the single insert corresponding to *3WJdB-T*, *Top-T*, and *Bottom-T* were cloned using standard restriction digest cloning to insert the sequence between the XbaI and XmaI sites in the pUC19 plasmid and were terminated with the T7 terminator. Dual expression plasmids contained the T7 terminator immediately downstream of *Top* and the T500 terminator derivative immediately downstream of *Bottom*. DNA assembly (NEBuilder HiFi DNA Assembly, New England Biolabs) was used to insert the two inserts, along with a 270-nucleotide spacer sequence, between the XbaI and XmaI sites of the pUC19 plasmid. All plasmid inserts were confirmed through sequencing. Plasmids for *in vivo* transcription by native *E. coli* RNA polymerase were deposited to Addgene (IDs 87311-87315). For details of DNA sequences refer to Supporting Information.

The protocol for *in vivo* assessment in *E. coli* was adapted from the Spinach and Broccoli publications.^17,28^ In short, 100 ng of each plasmid was transformed into *E. coli* BL21(DE3) competent cells using a standard heat shock transformation protocol. Cells were plated on 2xYT-agar plates containing 100 μg/mL of ampicillin and incubated overnight at 37°C. Single colonies from each plate were cultured overnight (∼16 hr) at 37°C with 250 RPM shaking in 3 mL of 2xYT broth containing 100 μg/mL of ampicillin. Following overnight culture, 100 μL of samples were added to 2.9 mL of fresh liquid media until reaching OD_600_ = 0.4, at which point 500 μL of sample was pelleted at 2,000 RCF and resuspended in 1 mL Dulbecco’s phosphate-buffered saline (DPBS) with calcium and magnesium (Thermo Fisher). 200 μL aliquots were then incubated with 200 μM DFHBI-1T at 37°C for 45 min prior to assessment on an Accuri C6 flow cytometer (BD Biosciences) configured with a 488 nm excitation laser and 533/30 nm filter. For each population, 5 x 10^4^ events were analyzed and processed using FlowJo software.

Samples for fluorescence microscopy were prepared as described above. Following the centrifugation of freshly diluted overnight culture, samples were resuspended in 200 μL of DPBS and plated on Poly-D-lysine-coated 8-chamber glass chamber slides (Lab-Tek). After a 45 min incubation at 37°C, cells were washed twice in DPBS and incubated with 200 μM DFHBI-1T in DPBS for an additional 45 min. Fluorescence images were obtained at 100X oil objective magnification using QCapture Suite Plus software and Rolera-XR camera (QImaging) mounted on an Olympus 1X70 inverted fluorescent microscope. Green fluorescence (λ_ex_ = 470/40 nm, λ_em_ = 525/50, 33 ms exposure) and bright field images were captured and exported into Fiji^53^ for linear adjustment of the brightness and contrast.

### Split-Broccoli Toehold Switch Assays

Toehold (TS2_KS01) and Trigger (TS2_AT01) RNA sequences were used as published.^9^ Inserts for *Top-Toehold-mCherry* and *Trigger-Bottom* were synthesized as gBlocks gene fragments from Integrated DNA Technologies and cloned into pUC19 between the XbaI and XmaI restriction sites. For the dual-insert plasmid containing *Top-Toehold-mCherry* and *Trigger-Bottom* (“Split-Broccoli Toehold Switch”), the pUC19-Trigger-Bottom plasmid was linearized with NdeI and the insert for *Top-Toehold-mCherry* was inserted using Gibson assembly. Each transcription unit contained a constitutive *E. coli* P70a promoter^52^ and a T7 transcription terminator. *In vivo* analysis was performed as described above for *in vivo* fluorescence, with additional detection for red fluorescence using a 670 nm low-pass filter. For each population, 5 x 104 events were analyzed and processed using FlowJo software. Fluorescence microscopy was performed as described for *In Vivo Fluorescence,* with the addition of a red fluorescence filter set (λ_ex_ = 562/40 nm, λem = 624/40, 100 ms exposure).

TX-TL transcription and translation reactions (a gift of Julius B. Lucks) were prepared and performed as described.^52^ In short, *E. coli* cytoplasmic extract, reaction buffer with metabolites and 20 μM DFHBI-1T was added to either 8 nM plasmid or H_2_O (blank) on ice, in a total volume of 10 μL. Samples were immediately transferred to a flat-bottom 384-well plate and measured on a Synergy H1 plate reader (BioTek) configured with filter sets for green fluorescence (λ_ex_ = 472, λ_em_ = 507) and red fluorescence (λ_ex_ = 587, λ_em_ = 615 nm). Readings were taken at 3 min intervals over the course of 12 hours. Average blank values at each time point were subtracted from sample values before calculating mean and standard deviation. Plasmids and their sequences were deposited to Addgene (IDs 87314, 87316-87318). For details of DNA sequences refer to Supporting Information.

## Associated Content

### Supporting Information

Table S1, Figures S1-S10, Plasmid DNA Sequences and Annotated Maps.

## Author Information

### Corresponding Author

Twitter: @DHBurkeAguero, @BurkeLabRNA

## Note

The authors declare no competing financial interest.

## Author Contributions

K.K.A. and D.H.B. conceived the Split-Broccoli system. K.K.A., K.D.T., M.F.L., D.P. and D.H.B. designed the experiments. K.K.A., K.D.T., M.F.L. and D.P. performed the experiments. K.K.A. and D.H.B. wrote the manuscript.

## Acknowledgements

We acknowledge support for this work from the University of Missouri Department of Biochemistry, the National Institute of Allergy and Infectious Diseases of the National Institutes of Health (R01AI074389) and the National Aeronautics and Space Administration (NNX12AD66G). We thank members of the Burke lab, Julius B. Lucks (Northwestern University), Maureen McKeague (ETH Zurich) and Raghav R. Poudyal (Pennsylvania State University) for providing feedback on the manuscript.

## Supporting Information

**Table S1.**
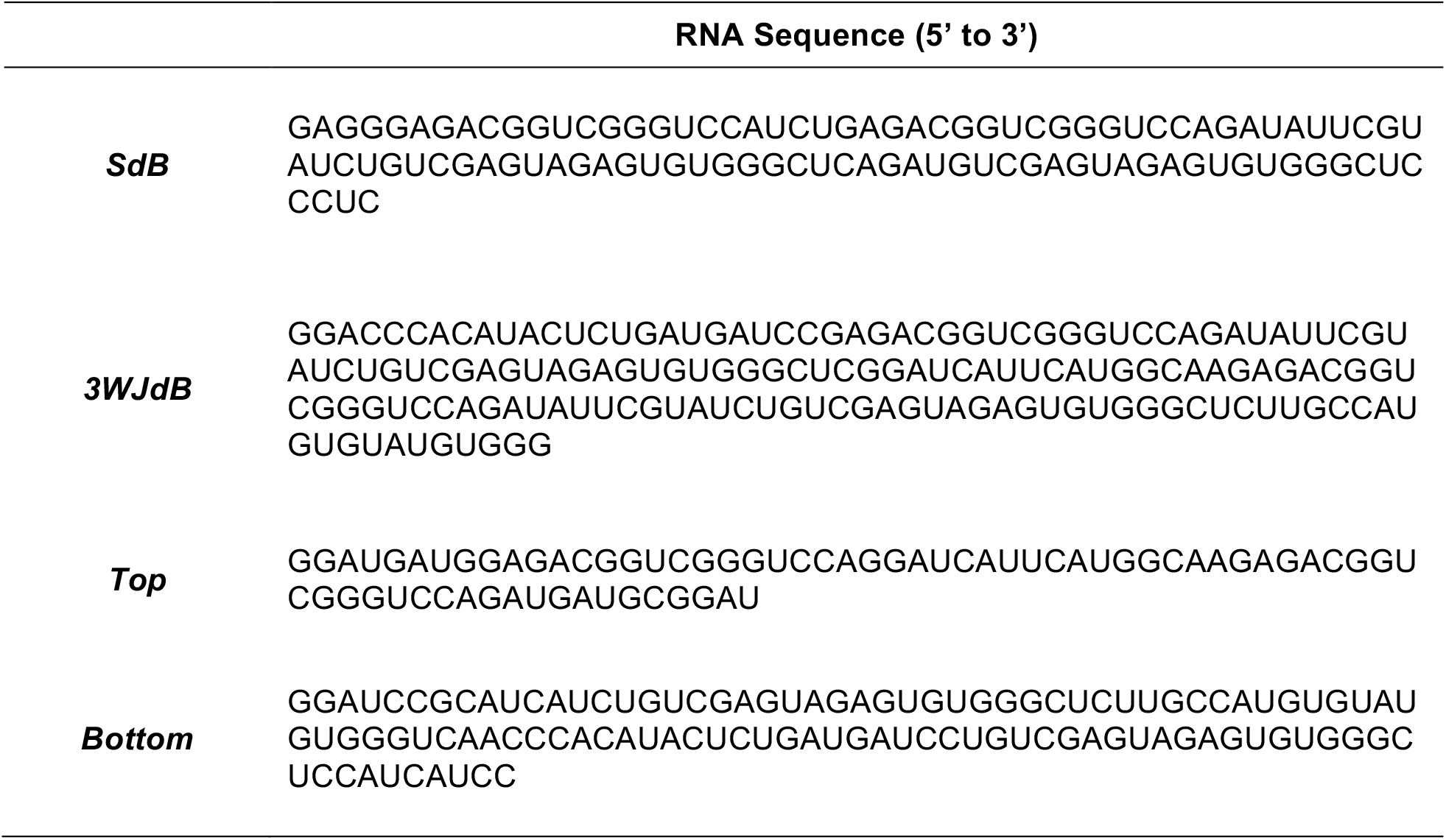
RNA sequences unique to this study.

**Figure S1.**
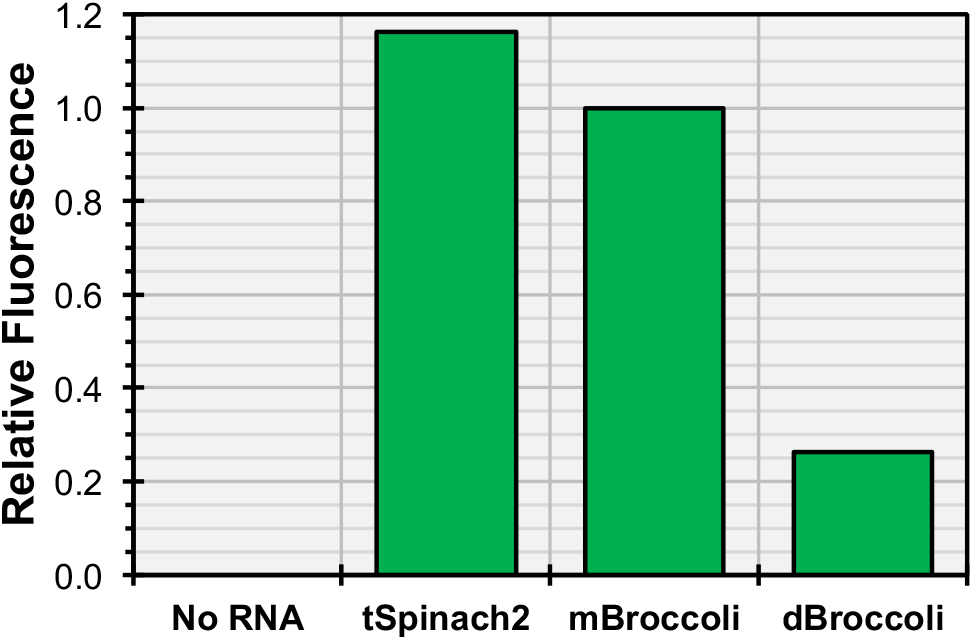
Comparison of *tSpinach2*, *mBroccoli* and *dBroccoli*. The Spinach2 aptamer embedded within a stabilizing tRNA^Lys^_3_ scaffold (*tSpinach2*) and the minimal monomeric Broccoli aptamer (*mBroccoli*) demonstrate similar levels of fluorescence when equimolar amounts of RNA are thermally renatured in buffer containing dye and assayed for fluorescence. In contrast, the minimal dimeric Broccoli aptamer (*dBroccoli*), which is expected to have twice the fluorescence of *mBroccoli*, exhibits about one-quarter the signal of *mBroccoli*, an 8-fold reduction from expected fluorescence activity.

**Figure S2.**
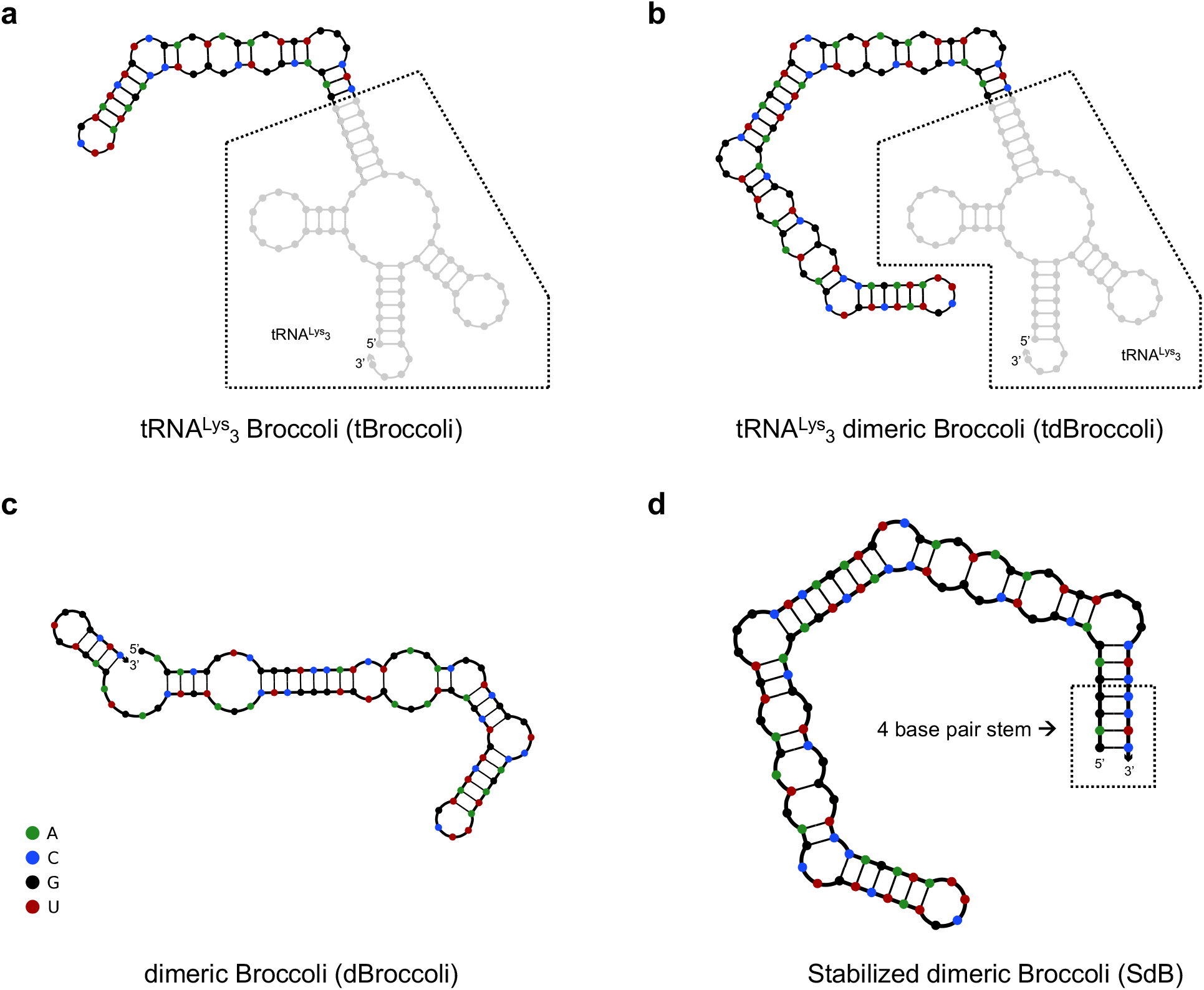
Predicted secondary structures of the Broccoli aptamer in the presence and absence of a tRNA stabilizing scaffold. (**a**) The Broccoli aptamer, when embedded within a tRNA^Lys^_3_ scaffold (*tBroccoli*), is predicted to form the functional fold of the Broccoli aptamer. (**b**) The dimeric Broccoli aptamer (*tdBroccoli*) is similarly stabilized within the tRNA scaffold. (**c**) In the absence of the scaffold, the minimal dimeric Broccoli (*dBroccoli*) is predicted to misfold by forming undesired Watson-Crick base pairs across each monomer that results in a nonfunctional aptamer. (**d**) Stabilized dimeric Broccoli (*SdB*) was designed by appending a 4 base pair terminal stem to the dimeric Broccoli and is predicted to form the functional aptamer fold in the absence of a larger scaffold.

**Figure S3.**
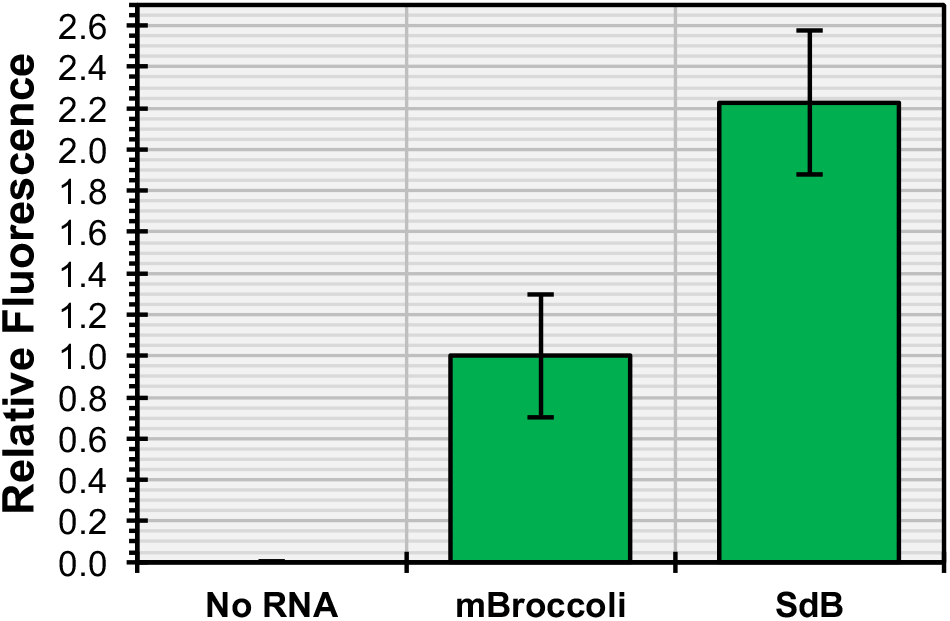
Stabilized dimeric Broccoli (*SdB*) is twice as bright as *mBroccoli*. *SdB* generates more than twice the signal of a monomeric Broccoli (*mBroccoli*) when equimolar amounts of *mBroccoli* and *SdB* aptamers are thermally renatured in buffer containing dye and assayed for fluorescence. Mean values are shown (n ≥ 6) with error bars to indicate standard deviation.

**Figure S4.**
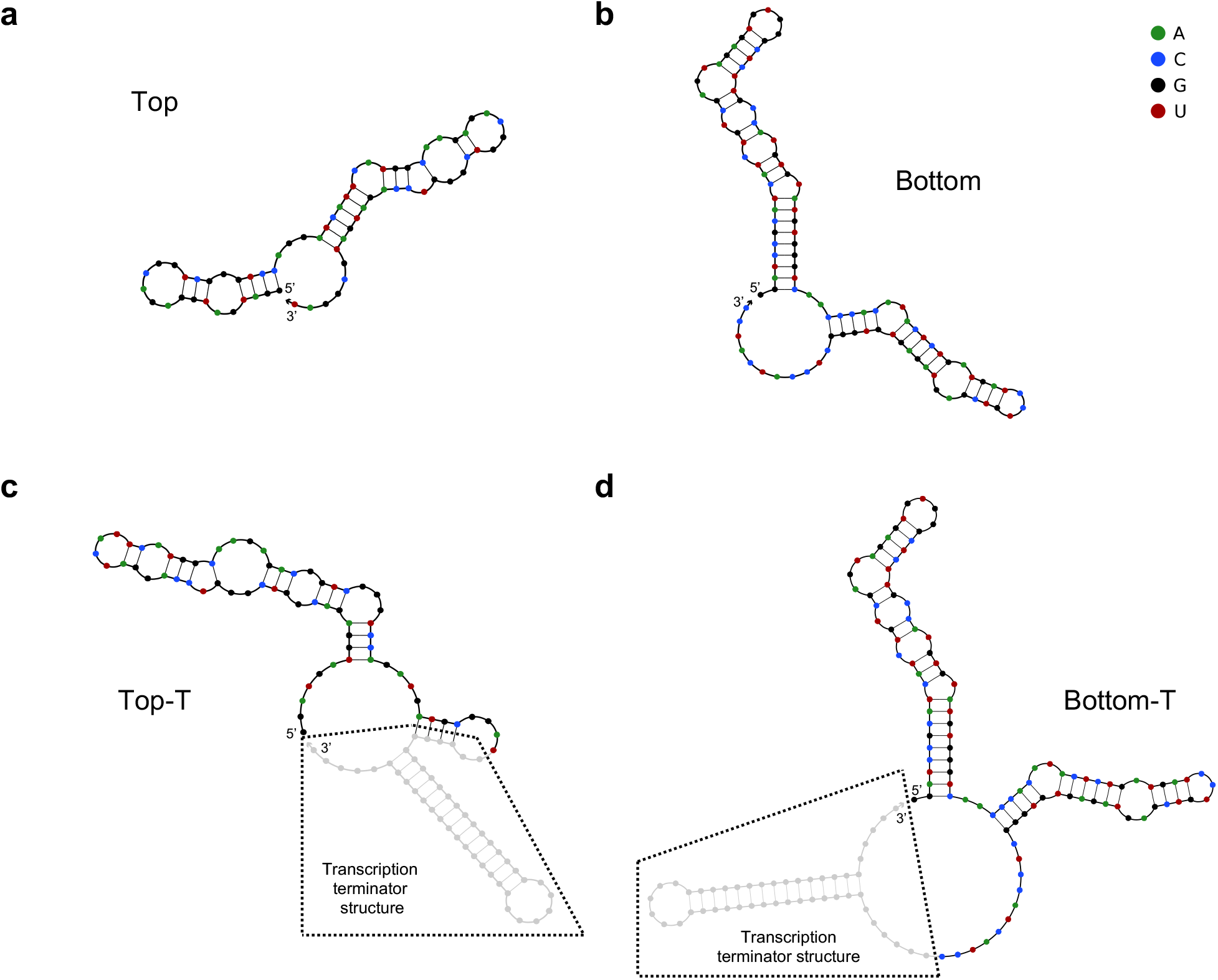
Predicted secondary structures of the Split-Broccoli system components, with and without transcription terminators. RNA corresponding to *Top* (**a**) and *Bottom* (**b**), without transcription terminator structures, and *Top-T* (**c**) and *Bottom-T* (**d**), which contain transcription terminator structures (boxed in lower panels). All four sequences neither contain the full Broccoli aptamer sequence, nor are independently predicted to fold into structures that resemble the secondary structure of the Broccoli aptamer.

**Figure S5.**
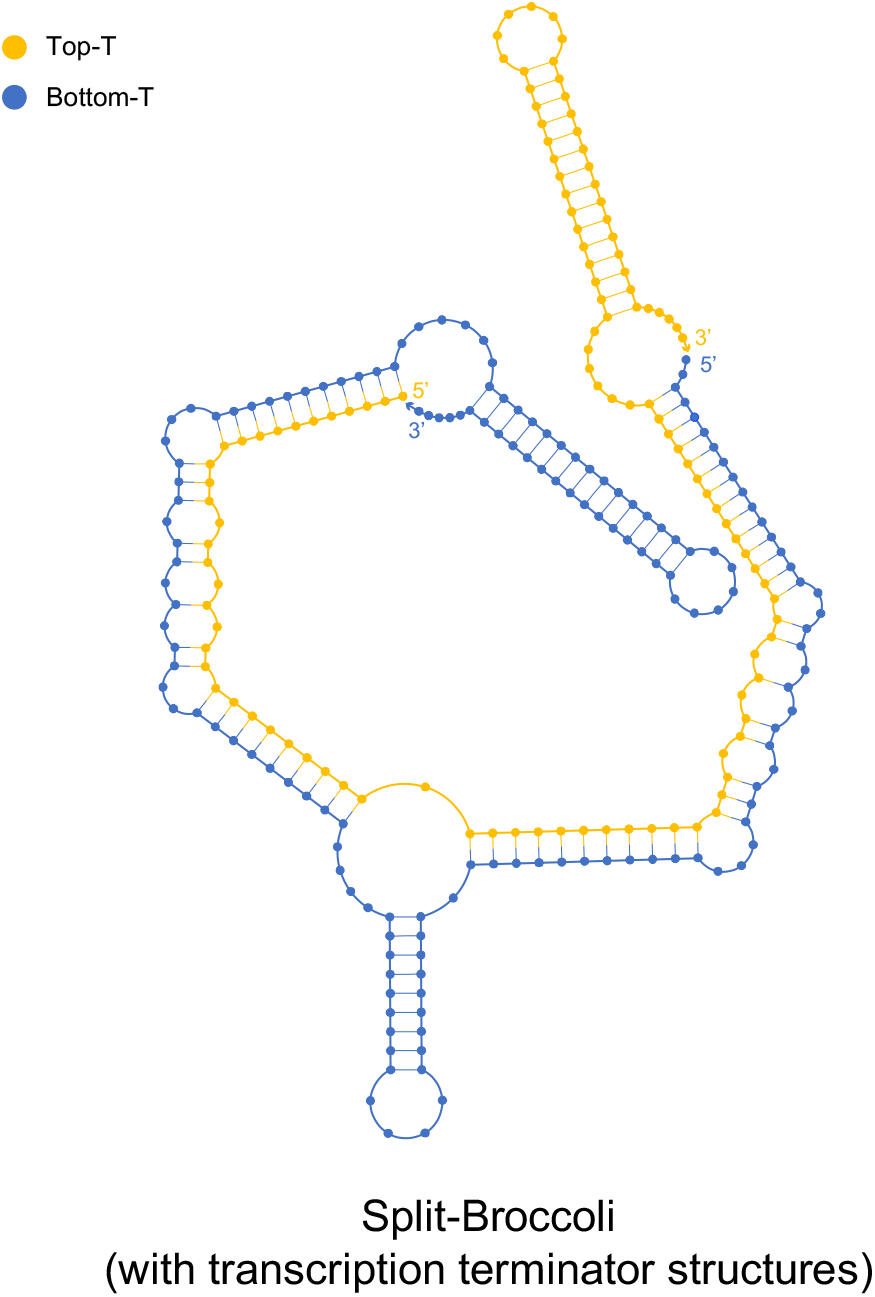
Predicted secondary structure of the hybridized Split-Broccoli system with transcription terminators. *Top-T* (yellow) and *Bottom-T* (blue) are predicted to hybridize into a secondary structure that maintains the expected motifs corresponding to the 3WJ, the Broccoli aptamers and the transcription terminators.

**Figure S6.**
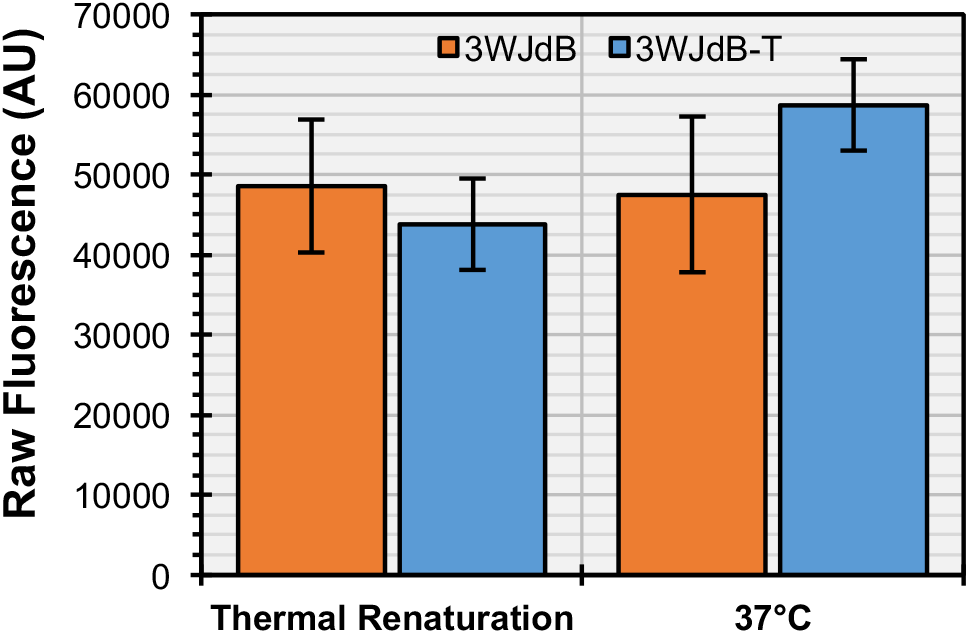
Comparison of raw fluorescence values for *3WJdB* with and without a transcription terminator, and with and without a thermal renaturation step. *3WJdB* without (orange columns) and with a transcription terminator structure (*3WJdB-T*, blue columns) demonstrate similar levels of fluorescence when assembled and measured (λ_ex_ = 472 nm, λ_em_ = 507 nm) following thermal renaturation from 90°C to 37°C (left columns) or simple incubation at 37°C (right columns). Mean values are shown (n = 5) with error bars to indicate standard deviation.

**Figure S7.**
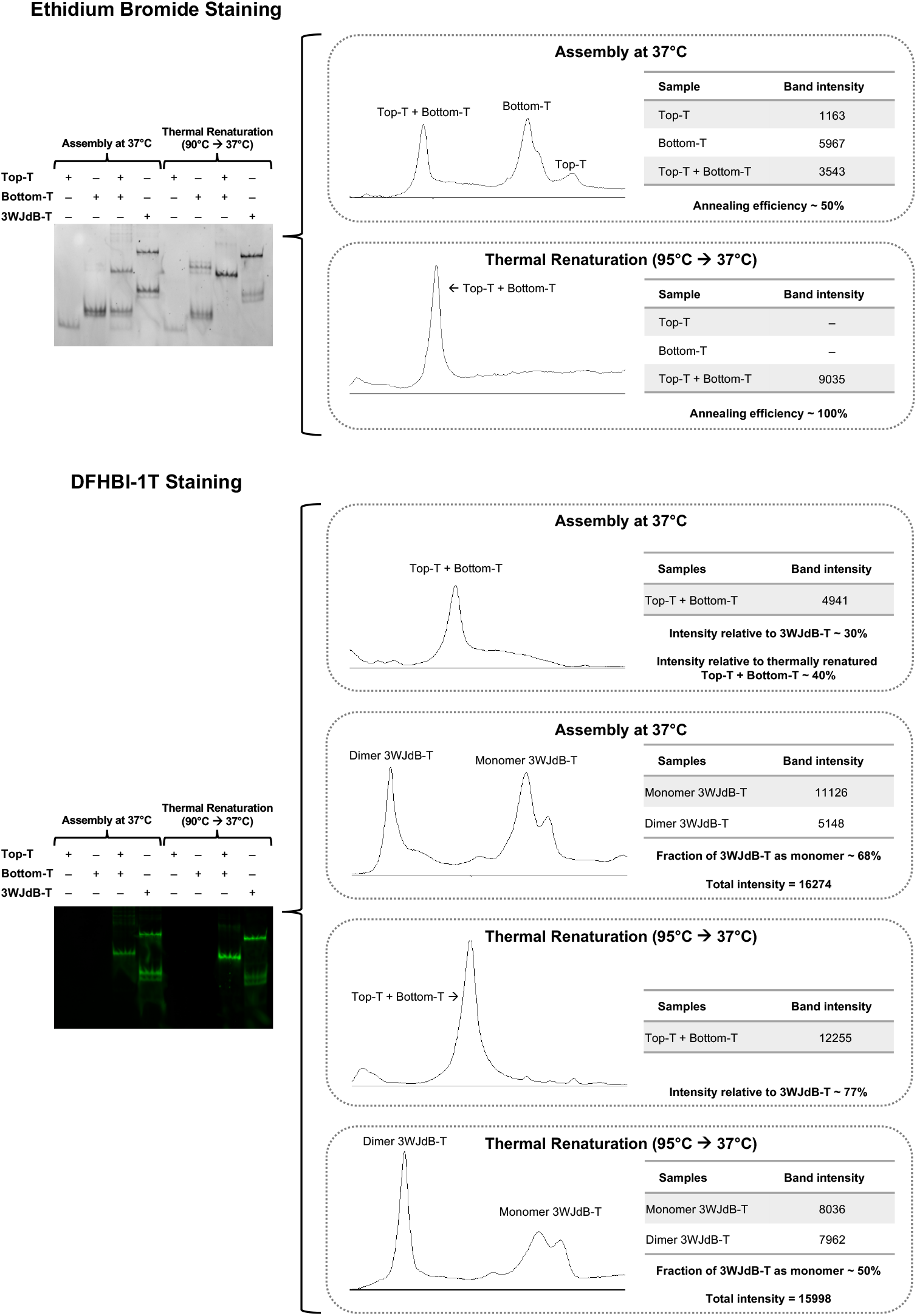
Densitometry analysis of the Split-Broccoli system with transcription terminators. Ethidium bromide staining identifies all molecular species (top). While DFHBI-1T staining (bottom) identifies only the functional, fluorescence-activating species. Thermal renaturation improves both annealing efficiency and fluorescence of the Split-Broccoli system.

**Figure S8.**
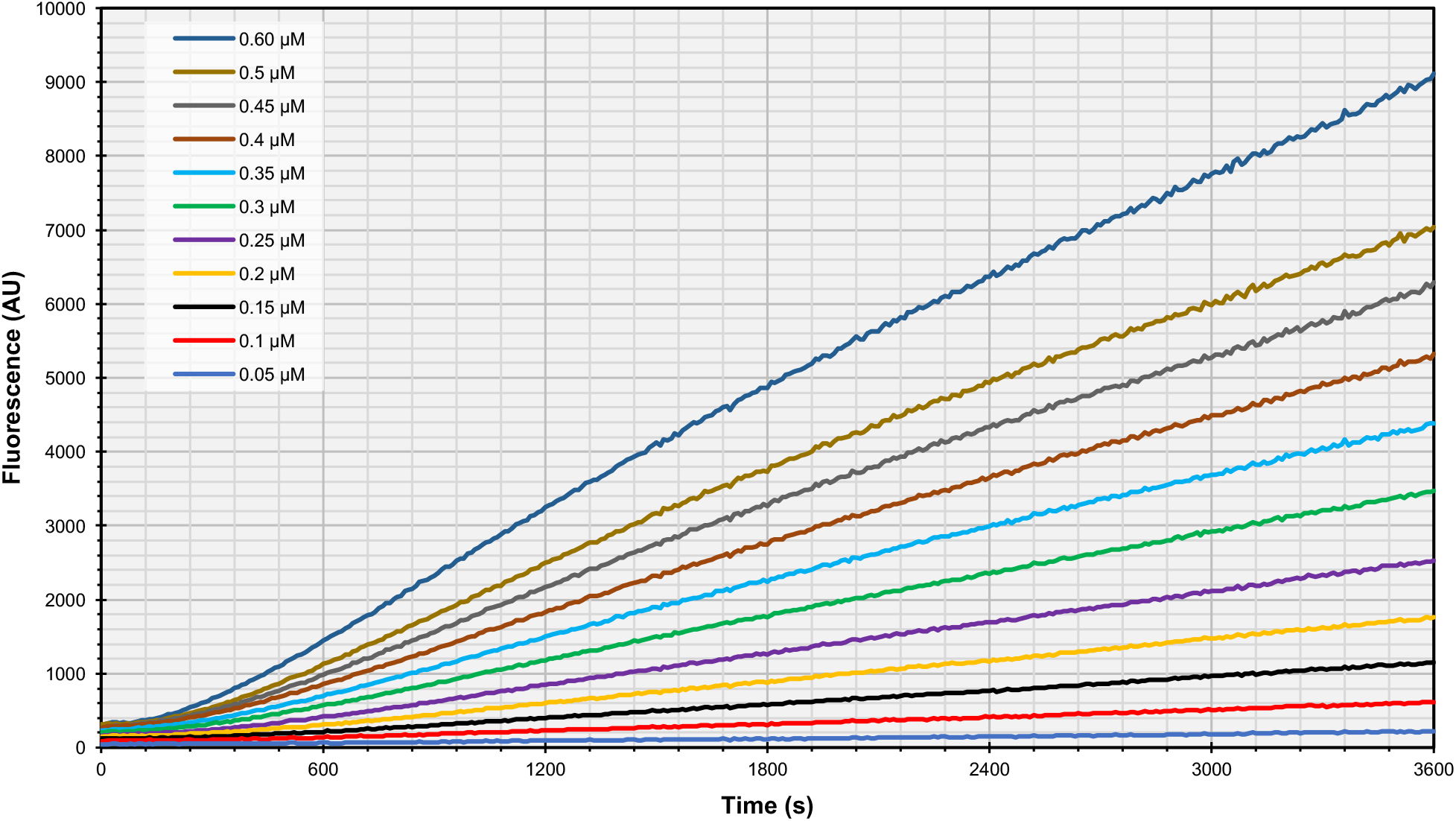
Fluorescence activation kinetics of the Split-Broccoli system with transcription terminators at varying concentrations. Functional assembly of equimolar amounts of *Top-T* and *Bottom-T* at 37°C was assayed at a range of concentrations. Rates from the linear region of each concentration replicate were then used to determine the rate of assembly. Mean values (n = 4) are shown for each concentration.

**Figure S9.**
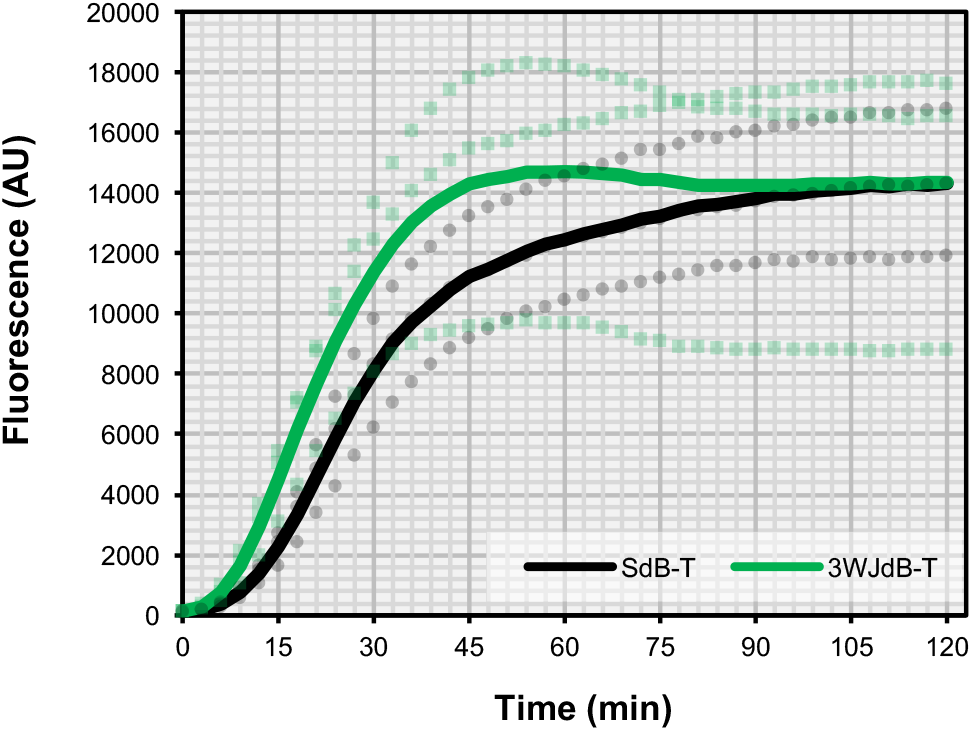
Addition of the 3WJ scaffold does not impede function of dimeric Broccoli when transcribed and assayed for activity *in vitro*. Transcriptional assays of *3WJdB-T* (green lines) and *SdB-T* (black lines) demonstrate that the addition of the 3WJ scaffold present in *3WJdB-T* does not slow the rate of fluorescence activation of dimeric Broccoli when compared to the unscaffolded variant, *SdB-T*. Mean values (n = 3) are shown as solid lines and individual replicates are shown as dotted lines.

**Figure S10.**
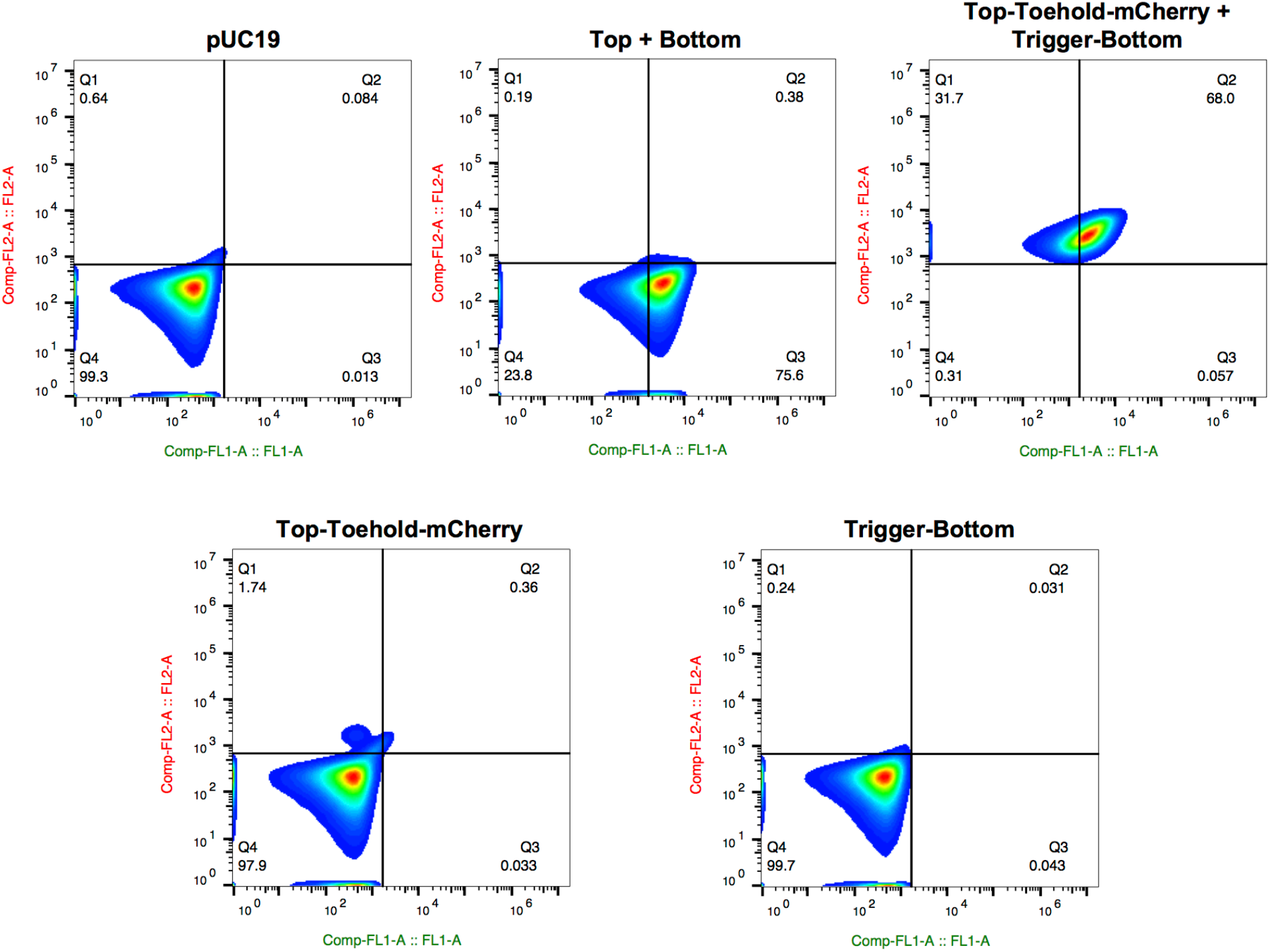
Representative scatter plots from flow cytometric analysis of the Split-Broccoli Toehold Switch demonstrate hybridization and activation of Split-Broccoli and the Toehold switch. *E. coli* containing a plasmid encoding the Split-Broccoli system fused to an RNA toehold switch (*Top-Toehold-mCherry* + *Trigger-Bottom*, top right) demonstrate a notable shift in both green and red fluorescence (x- and y-axes, respectively), while *E. coli* containing a plasmid encoding the Split-Broccoli system alone (*Top* + *Bottom*, top middle) only exhibits a shift in green fluorescence. In contrast, *E. coli* transformed with a plasmid encoding either *Top-Toehold-mCherry* (bottom left) or *Trigger-Bottom* alone (bottom right) do not cause a major shift in either red or green cellular fluorescence.

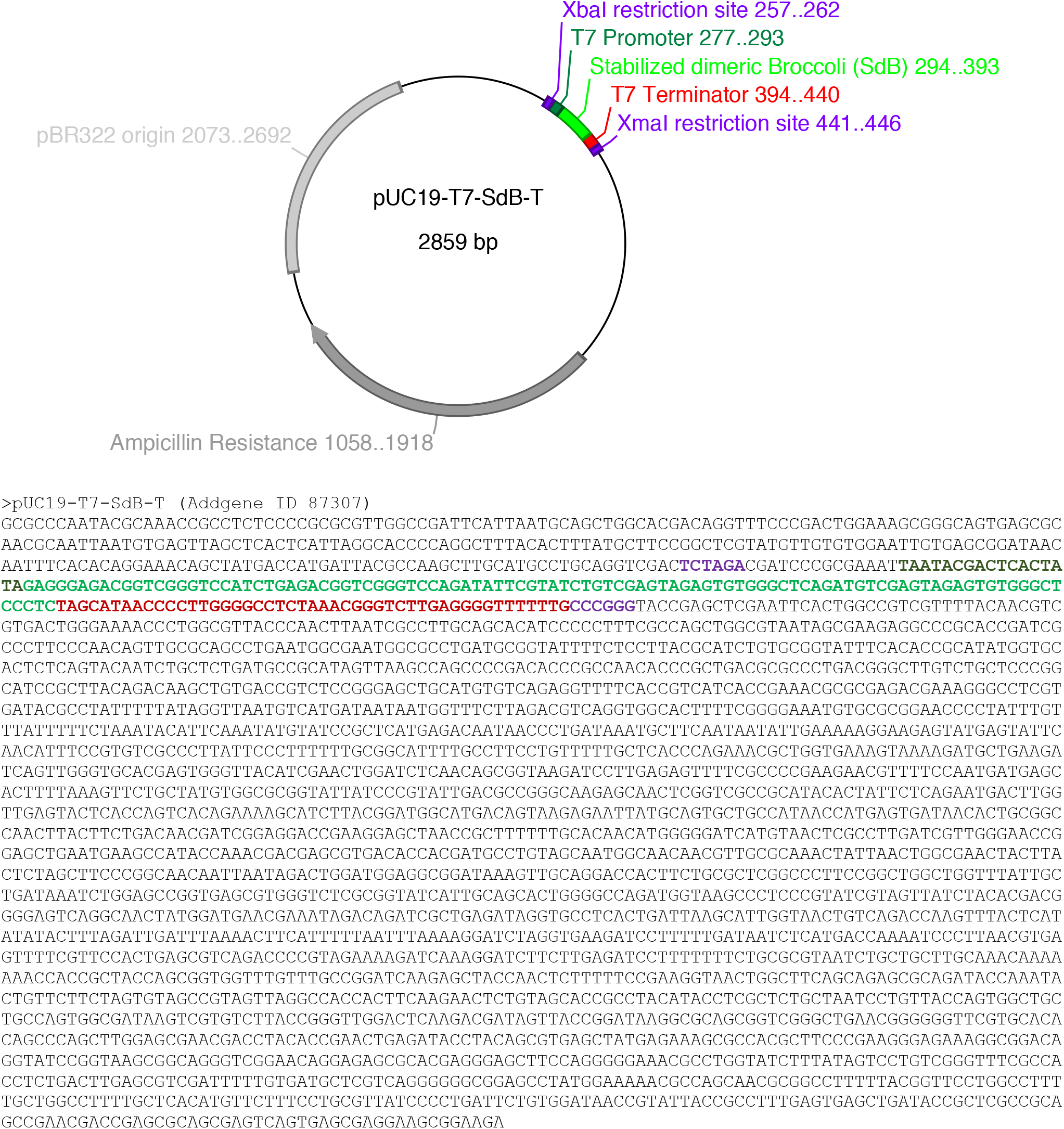

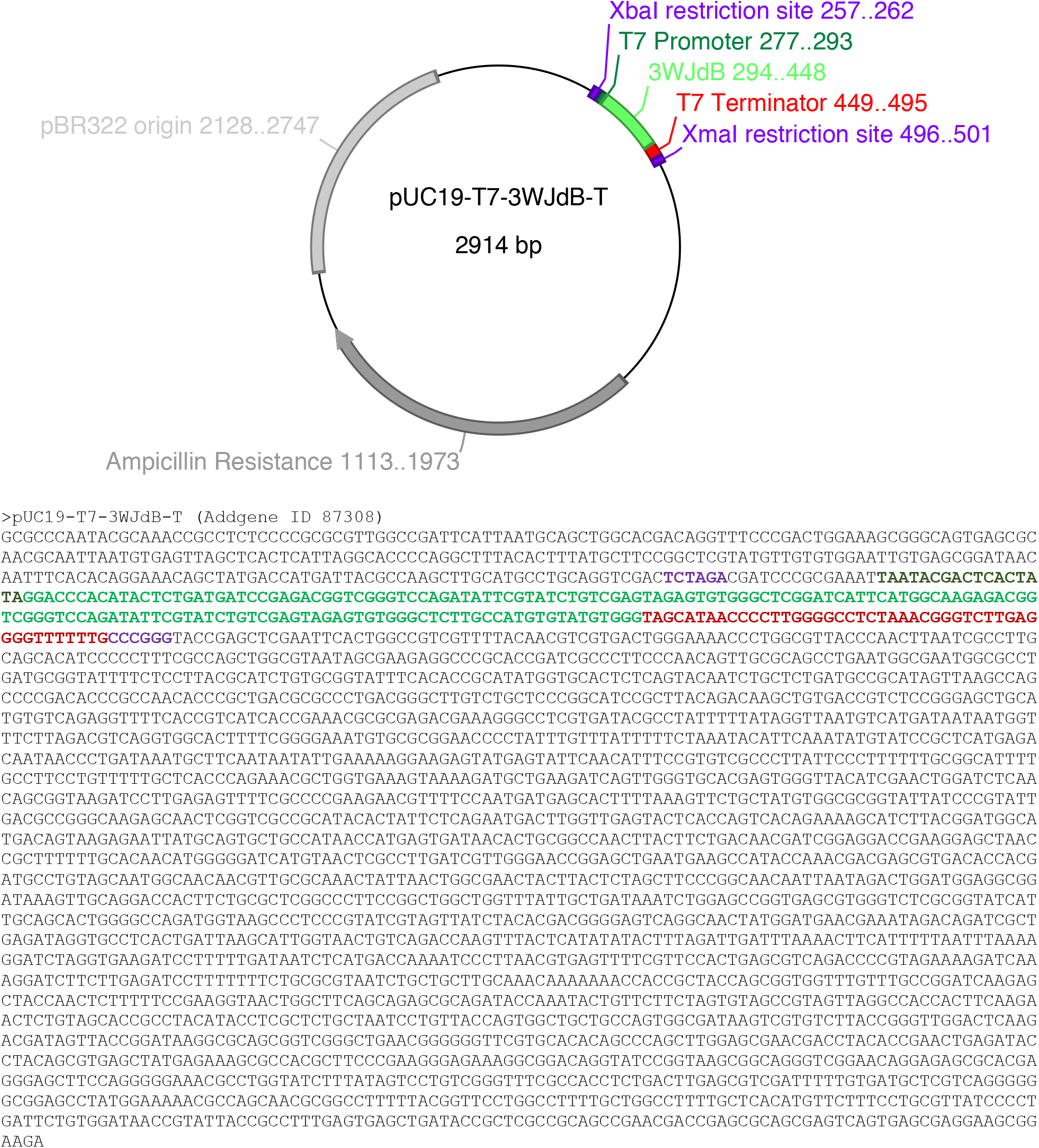

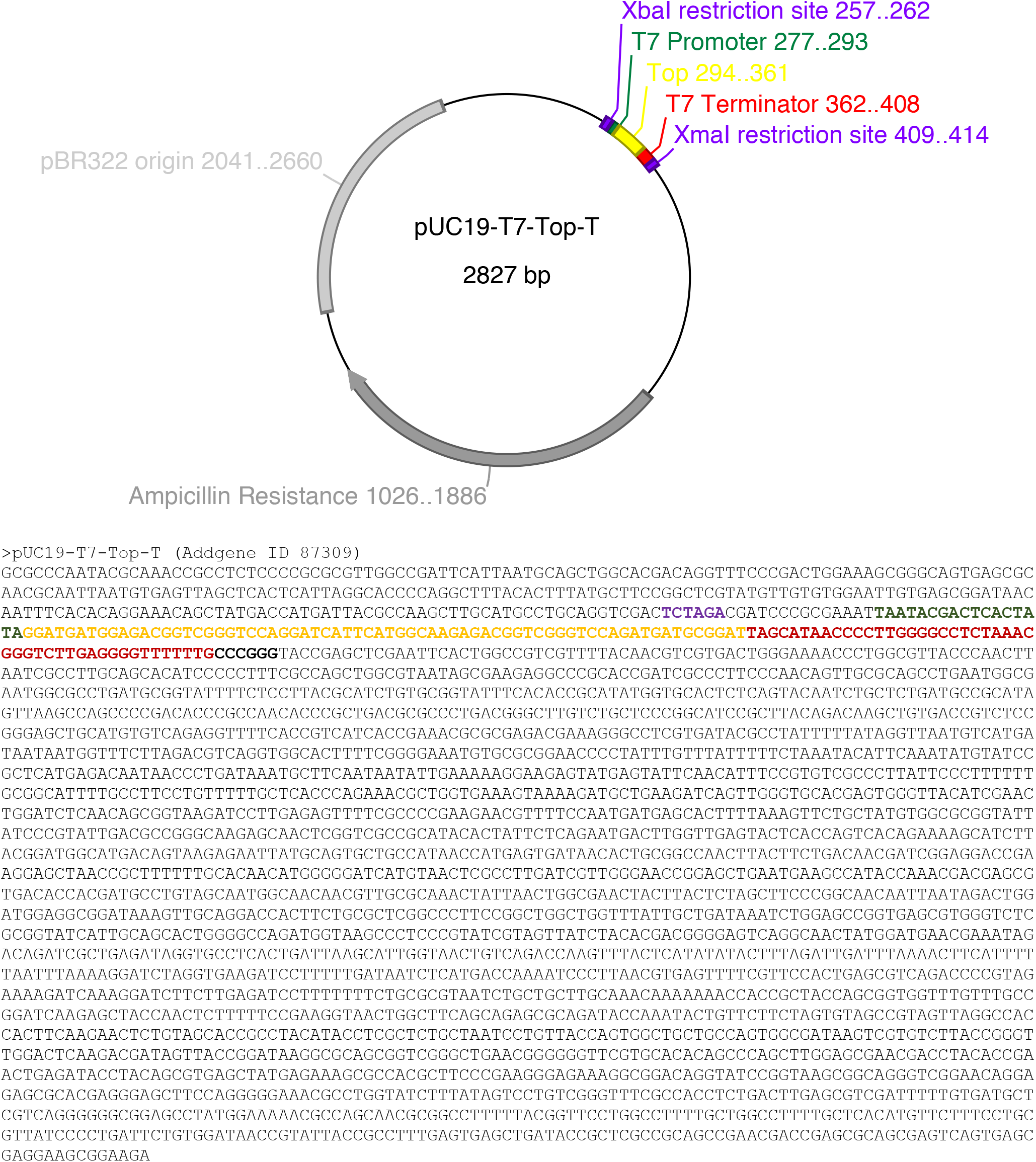

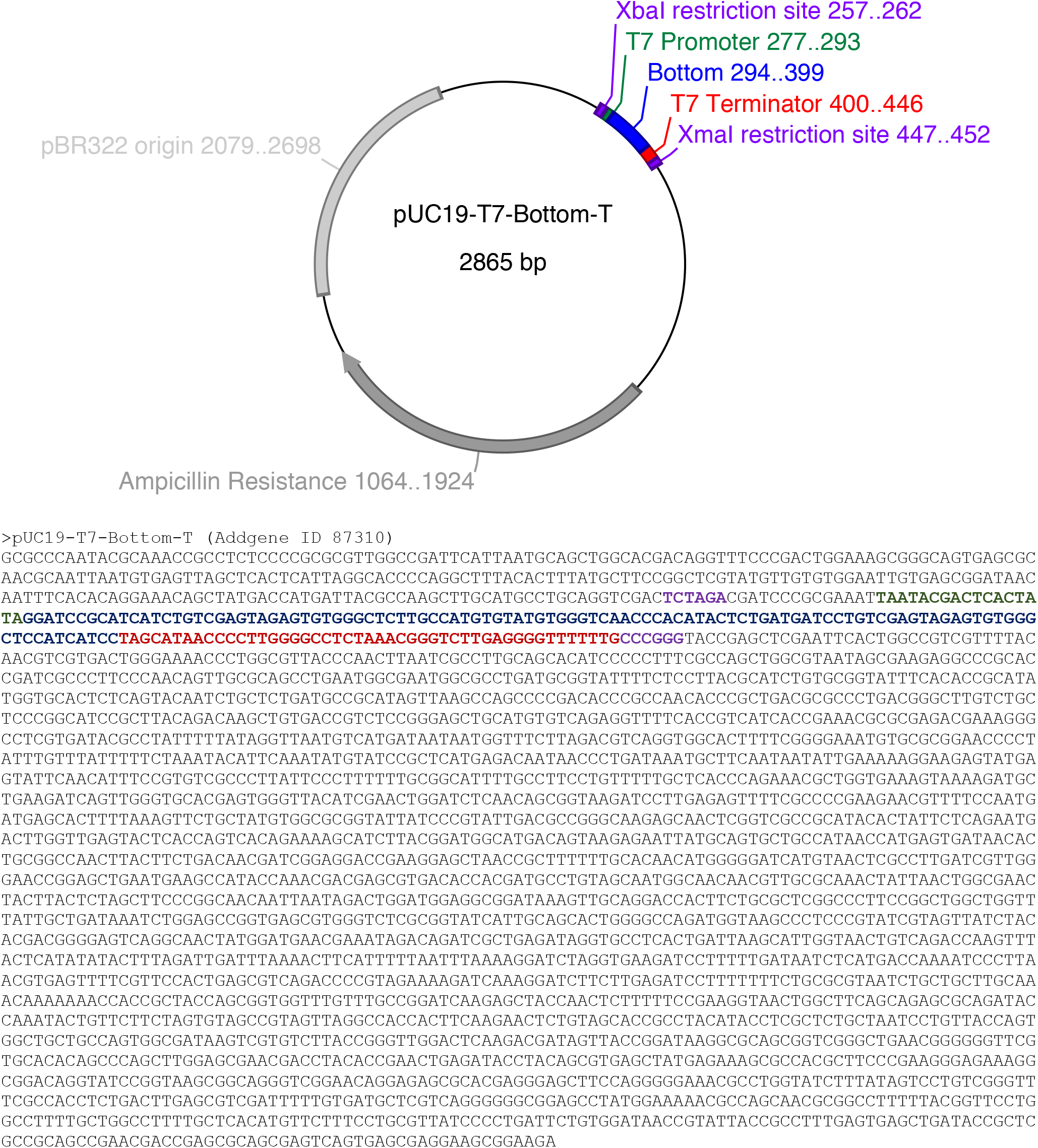

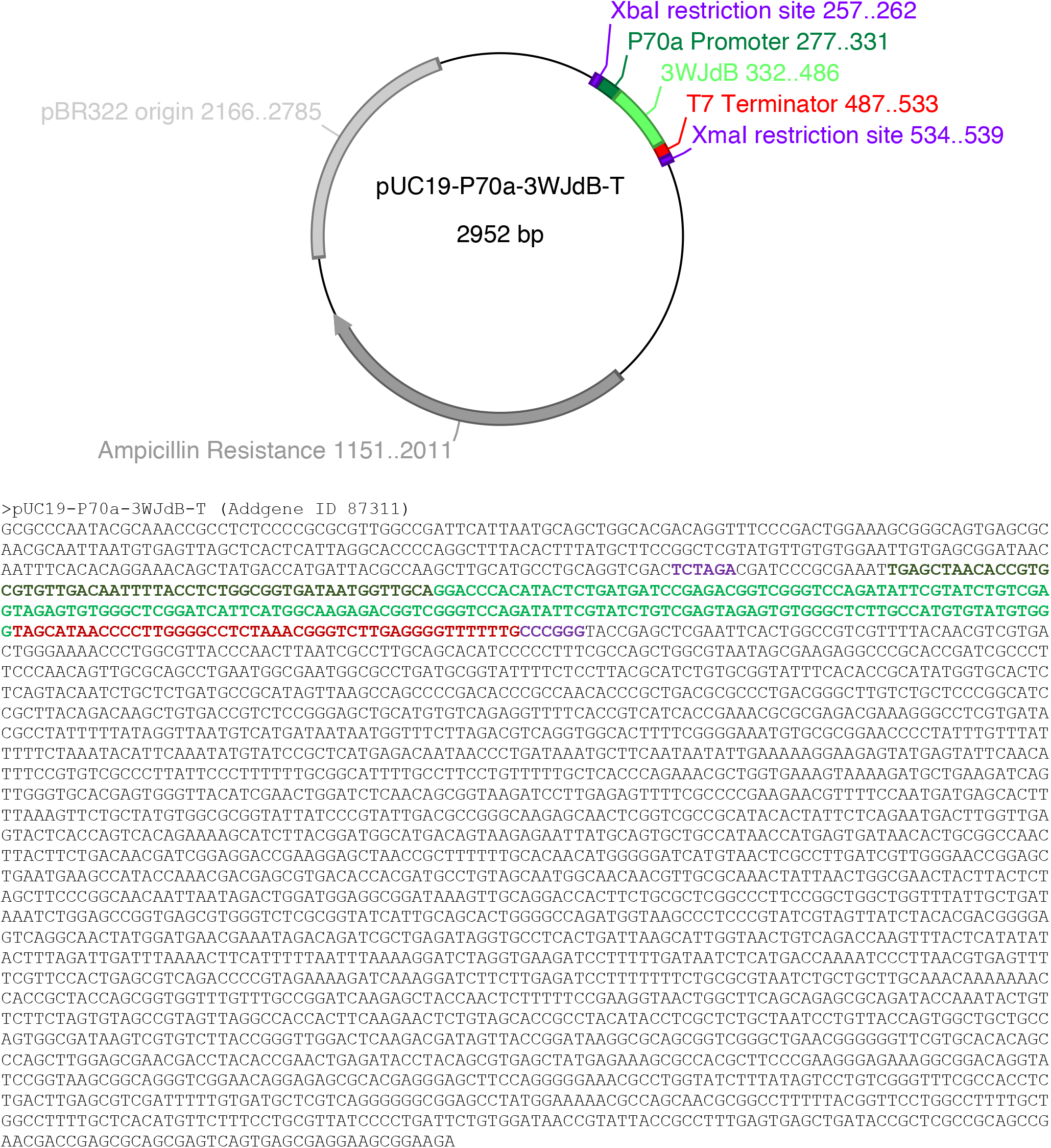

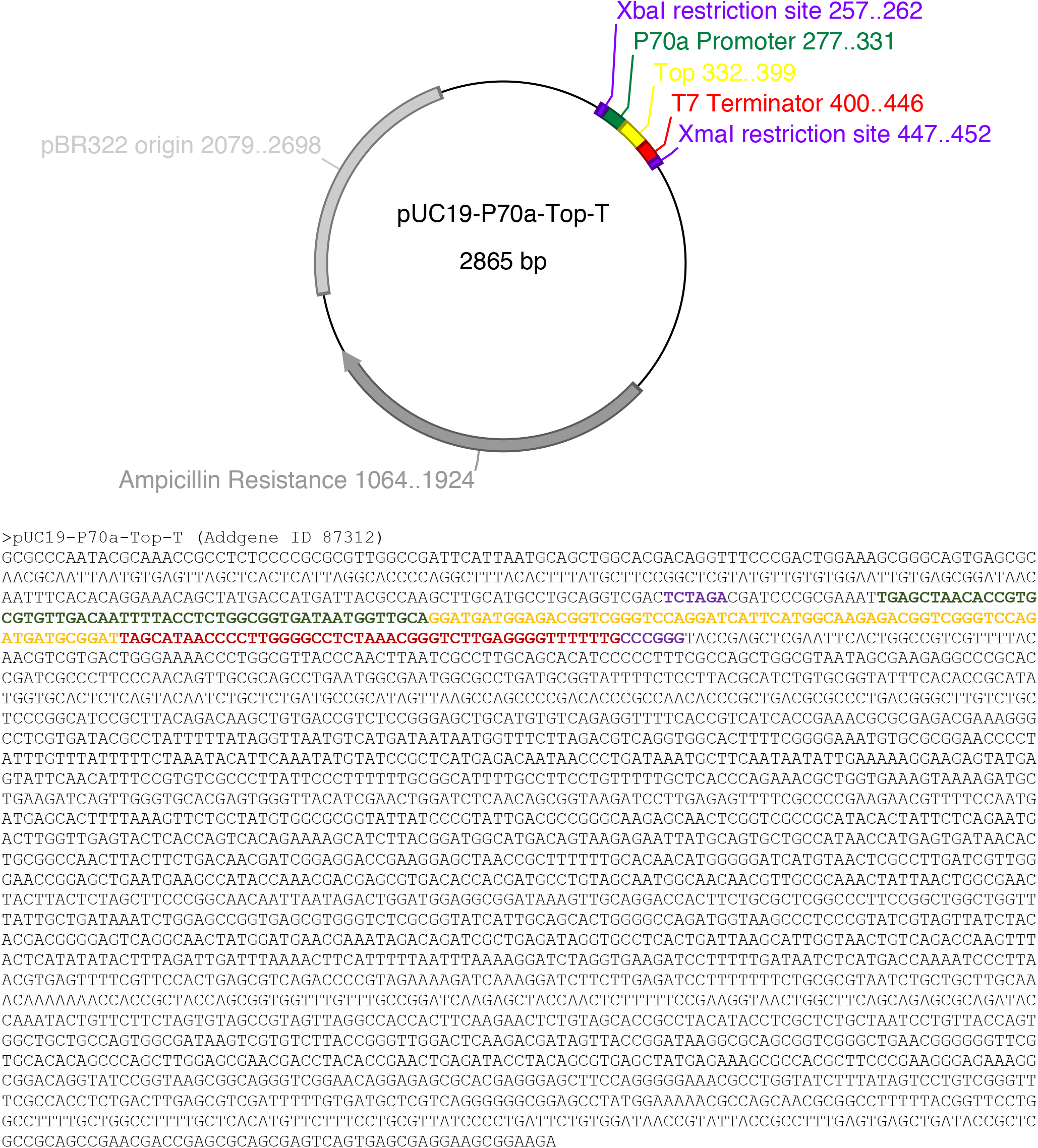

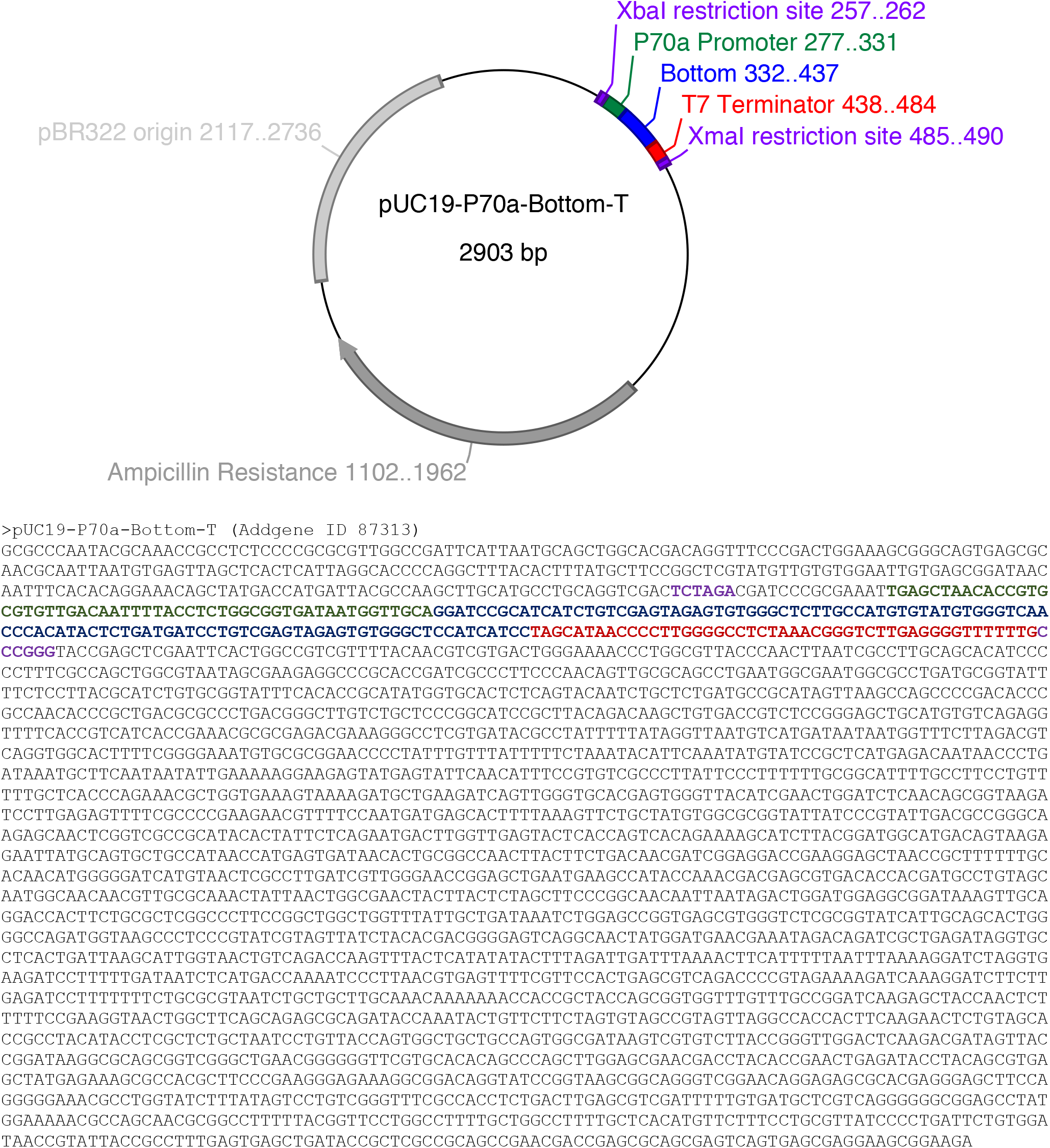

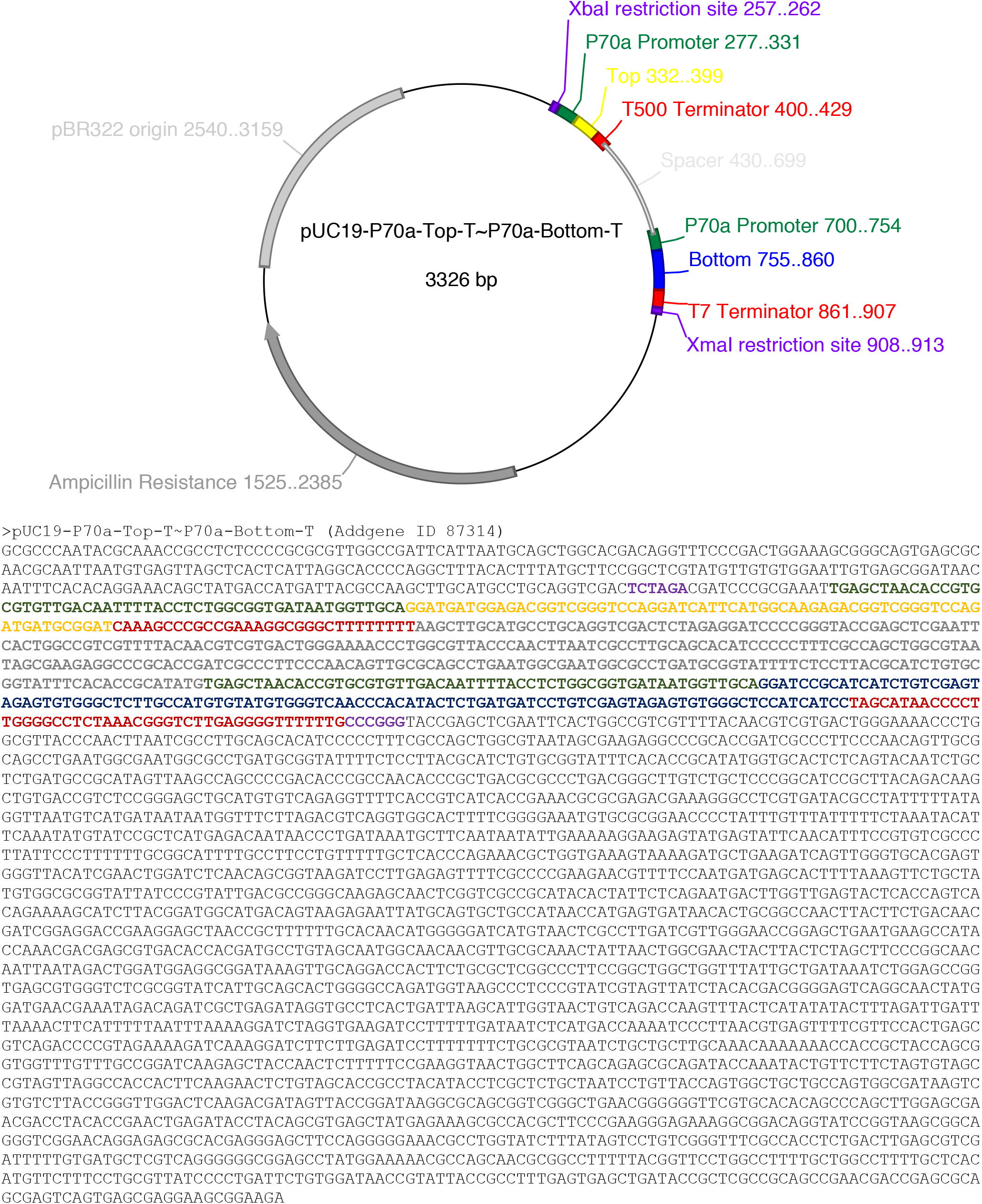

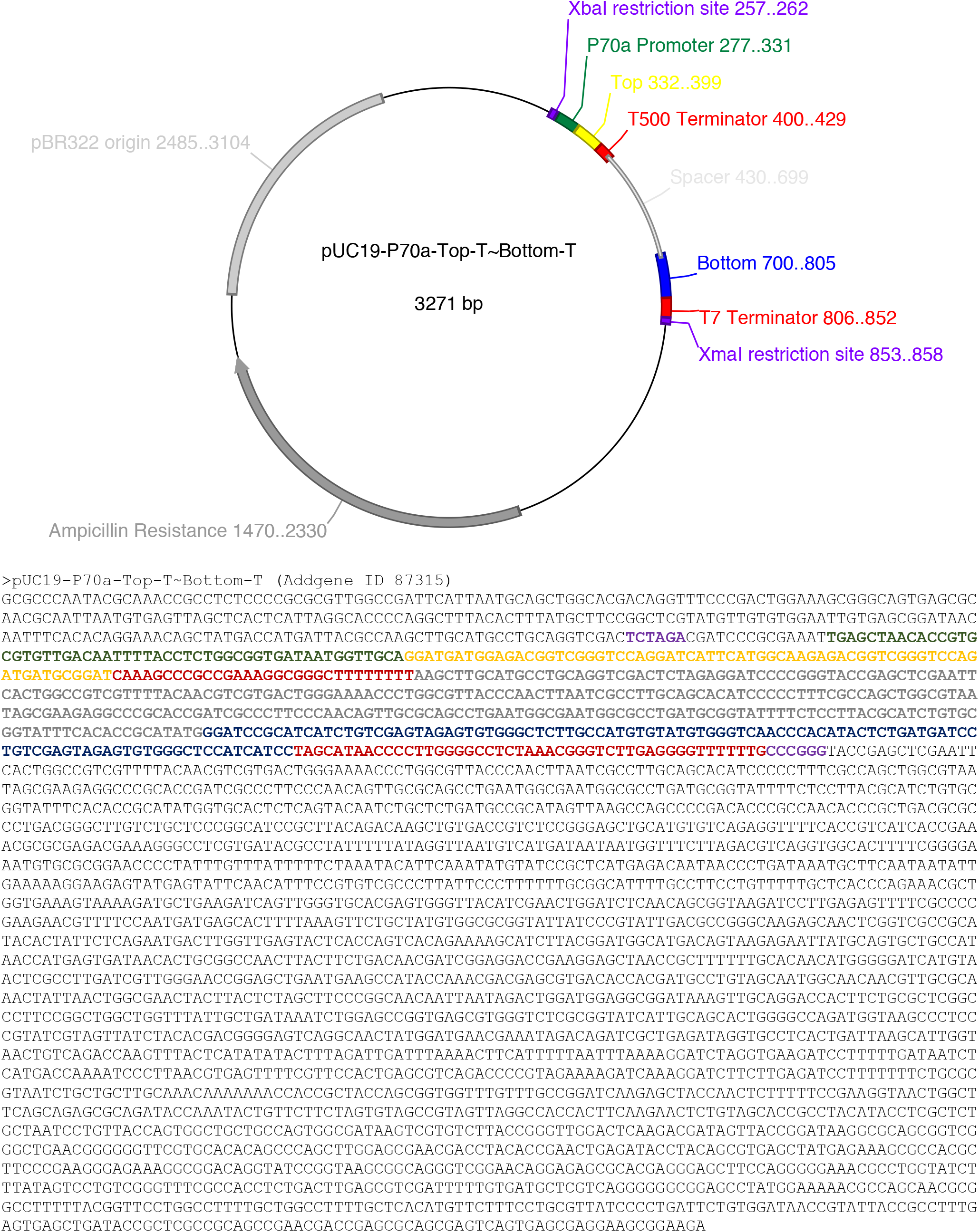

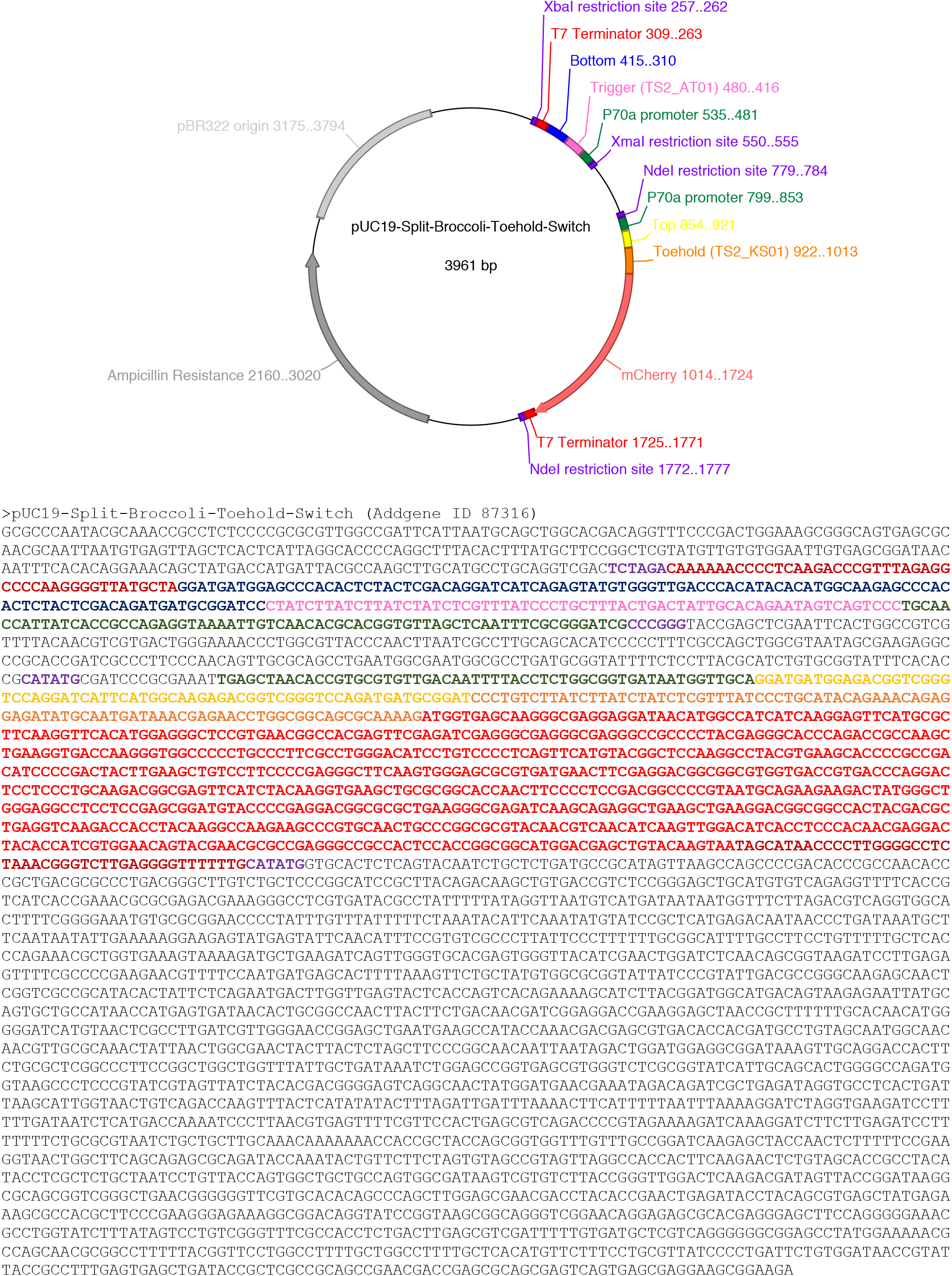

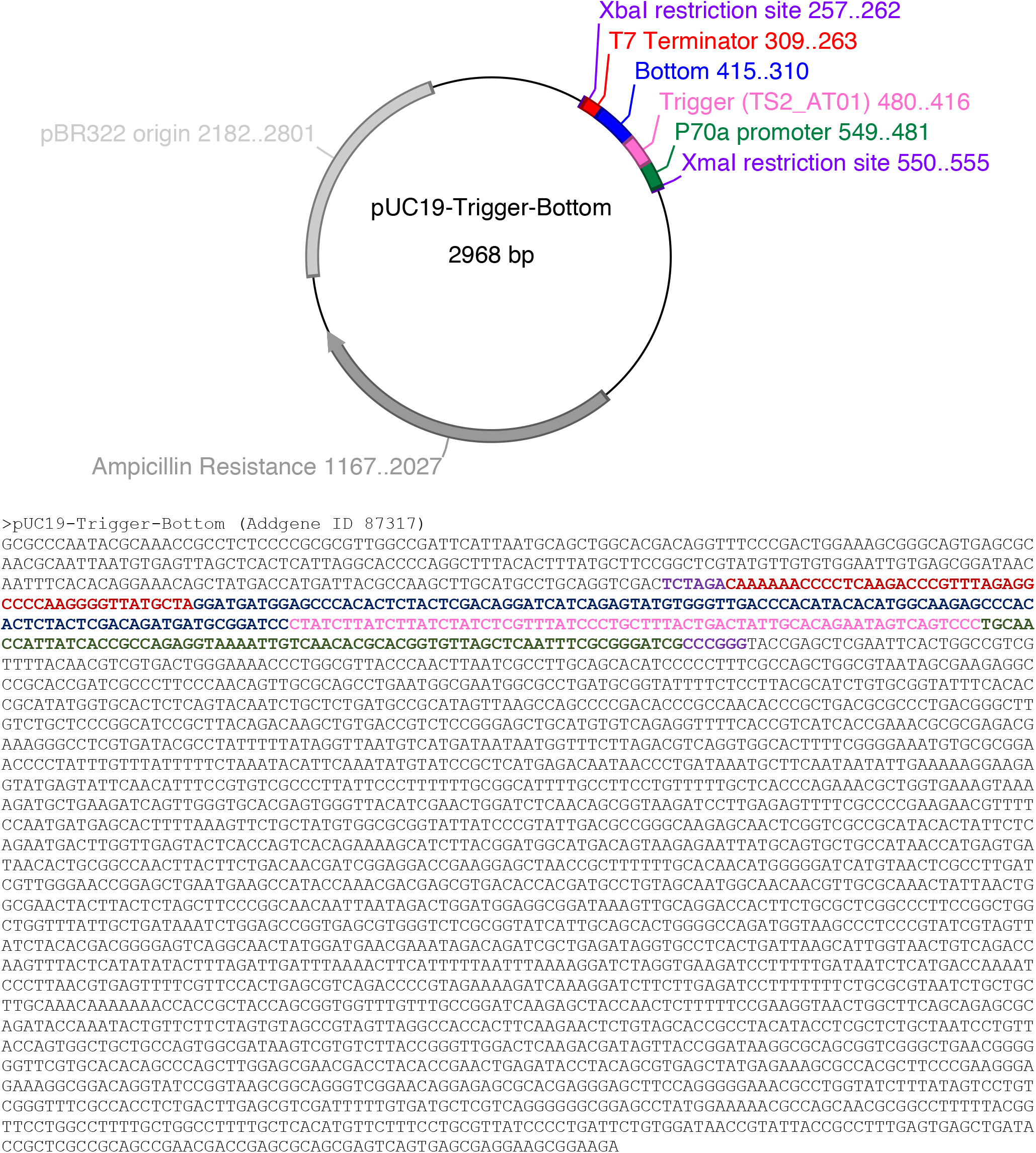

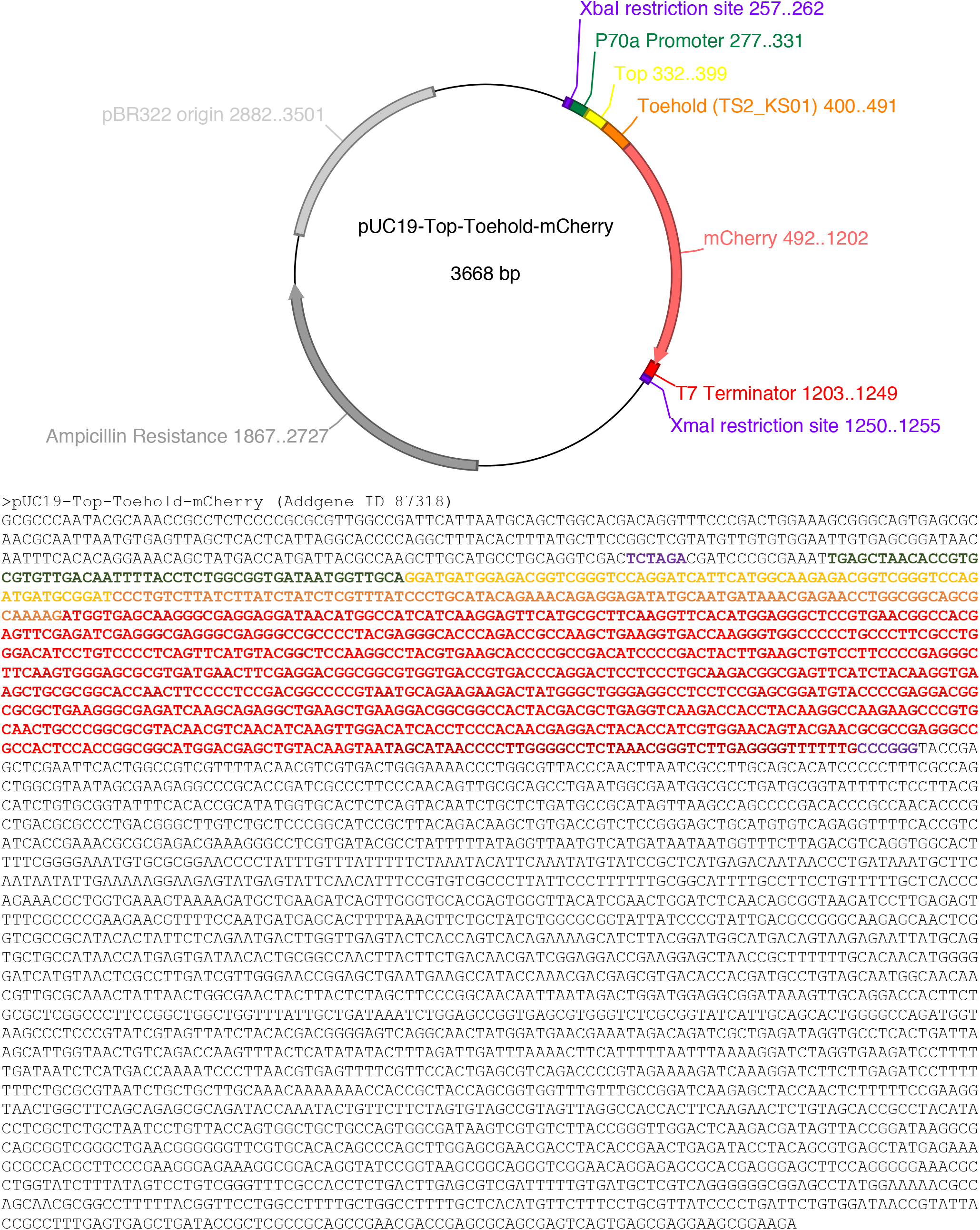

